# Frequency-dependence in multidimensional diffusion-relaxation correlation MRI of the brain: overfitting or meaningful parameter?

**DOI:** 10.1101/2024.04.29.591586

**Authors:** Maxime Yon, Omar Narvaez, Jan Martin, Hong Jiang, Diana Bernin, Eva Forssell-Aronsson, Frederik Laun, Alejandra Sierra, Daniel Topgaard

## Abstract

Time- or frequency-dependent (“restricted”) diffusion potentially provides useful information about cellular-scale structures in the brain but is challenging to interpret because of the intravoxel tissue heterogeneity. Frequency-dependence was recently incorporated in a multidimensional diffusion-relaxation correlation MRI framework relying on tensor-valued diffusion encoding at multiple frequencies, as well as variable echo and repetition times, to give model-free characterization of intravoxel heterogeneity by Monte Carlo data inversion into nonparametric distributions of frequency-dependent diffusion tensors and nuclear relaxation rates. While microimaging equipment with high-amplitude magnetic field gradients allows exploration of frequencies from tens to hundreds of Hz, clinical scanners with more moderate gradient capabilities are limited to a narrow frequency range, which may be insufficient to observe effects of restricted diffusion for brain tissues. We here investigate the effects of including or omitting frequency-dependence in the data inversion from isotropic and anisotropic liquids, excised tumor tissue, ex vivo mouse brain, and in vivo human brain. For microimaging measurements covering a wide frequency range, from 35 to 320 Hz at *b*-values over 4·10^9^ sm^−2^, the inclusion of frequency-dependence drastically reduces the fit residuals and avoids bias in the diffusion metrics for tumor and brain voxels with micrometer-scale structures. Conversely, for the case of in vivo human brain investigated in the narrow frequency range from 5 to 11 Hz at *b* = 3·10^9^ sm^−2^, analyses with and without inclusion of frequency-dependence yield similar fit residuals and diffusion metrics for all voxels. These results indicate that frequency-dependent inversion may be generally applied to diffusion-relaxation correlation MRI data with and without observable effects of restricted diffusion.

## Introduction

MRI provides information about the brain at length scales below the millimeter-scale resolution of the imaging voxels via diffusion metrics (Jones, 2010), reporting on the micrometer-scale organization of macromolecules and cellular membranes acting as barriers for the tissue water (Beaulieu, 2002; Topgaard, 2020), and nuclear relaxation rates (Tofts, 2003) sensitive to the local concentrations of paramagnetic species (Zimmerman, 1954; Laukien and Schlüter, 1956) and chemically exchangeable protons on proteins (Edzes and Samulski, 1975) and carbohydrates (Hills et al., 1989). While quantitative diffusion and relaxation MRI measurements have traditionally been performed separately, recent developments have enabled adaption of multidimensional diffusion-relaxation correlation NMR methods (Galvosas and Callaghan, 2010; Bernin and Topgaard, 2013; Song et al., 2016) to improve characterization of intravoxel heterogeneity in the brain (Benjamini and Basser, 2020; Tax, 2020; Slator et al., 2021). More widespread applications of these new methods in neuroscience studies rely on identification of the most informative acquisition dimensions, design of time-efficient measurement protocols to explore the multidimensional acquisition space, and development of data processing methods with optimal trade-offs between flexibility and risk of overfitting and overinterpretation.

The MRI signal is sensitized to translational motion on the micrometer length-scale and the millisecond time-scale by application of time-dependent magnetic field gradients. On this length scale, the translational motion in isotropic liquids such as water is fully captured by the self-diffusion coefficient *D* (Stejskal and Tanner, 1965; Stilbs, 1987; Morris and Johnson Jr, 1992), and the relevant acquisition dimension is the “*b*-value” (Le Bihan et al., 1986). For anisotropic materials, the diffusion is described with a tensor **D** (Jost, 1952) which can be determined by performing a series of measurements where the relative orientation between the gradient and the object is varied as demonstrated for clay (Boss and Stejskal, 1965), wood (MacGregor et al., 1983), and brain white matter (WM) (Moseley et al., 1991). Here, the relevant acquisition variable is the encoding tensor **b** (Basser et al., 1994), which can be parameterized in terms of its magnitude, anisotropy, asymmetry, and orientation (Eriksson et al., 2015). In porous rocks (Woessner, 1963; Latour et al., 1993), emulsions (Packer and Rees, 1972; Topgaard et al., 2002), and biological tissues (Cooper et al., 1974; Tanner, 1979; Latour et al., 1994), where the investigated liquid is enclosed in or hindered by micrometer-scale objects, the measurements yield an apparent diffusion coefficient (ADC), which depends on the details of the timing parameters of the motion-encoding gradient waveform—in particular its overall duration which constitutes an additional acquisition variable. The time-dependence of ADC is in the diffusion NMR literature often referred to as “restricted” diffusion (Woessner, 1963; Stejskal, 1965; Packer and Rees, 1972; Cooper et al., 1974) and can be further analyzed to extract the surface-to-volume ratio, pore size, and tortuosity of porous media (Latour et al., 1993; Latour et al., 1995). The waveform duration also determines if the diffusivities of exchanging proton populations can be estimated individually or only as an average (Kärger, 1969; Johnson Jr, 1993).

Conventional diffusion MRI applied to the in vivo human brain is often performed with pairs of magnetic field gradient pulses with durations of tens of milliseconds, corresponding to displacements of a few tens of micrometers. For many tissues, this displacement is much larger than the typical cell sizes and thus yields ADC values that are independent of the experimentally accessible minor variations of the diffusion time and contain aggregated information about local diffusivities, cellular and sub-cellular structures, and barrier properties of the cell membranes (Le Bihan et al., 1993; Clark et al., 2001; Nilsson et al., 2009). Trains of gradient pulse pairs can be used to widen the range over which diffusion is monitored towards shorter time-scales and distances (Tanner, 1979; Schachter et al., 2000; Stepišnik and Callaghan, 2000; Clark et al., 2001; Topgaard et al., 2002). While the individual pulse pairs give insufficient diffusion weighting, as quantified by the *b*-value, their effect is accumulated over the duration of the pulse train. The periodicity of such “oscillating gradient spin-echo” (OGSE) diffusion encoding lends itself to analysis with a powerful frequency-domain formalism building on tensor-valued diffusion spectra **D**(*ω*) defined as the Fourier transform of the velocity correlation function (Stepišnik, 1981; Callaghan and Stepišnik, 1995). In this case, the tensor-valued encoding spectrum **b**(*ω*) is a useful acquisition variable (Nielsen et al., 2018; Topgaard, 2019b; Lundell and Lasič, 2020; Jiang et al., 2023). In materials with pores having simple and uniform geometries, OGSE can be used to quantify the surface-to-volume ratio (Parsons et al., 2003; Parsons Jr et al., 2006; Reynaud et al., 2015) and pore sizes (Parsons Jr et al., 2006; Li et al., 2014). Even without extracting quantitative geometrical information, OGSE is useful for providing contrast not available with conventional diffusion methods and has been applied in preclinical MRI at encoding frequencies up to 1 kHz (Portnoy et al., 2013) to highlight specific brain regions, such as the cerebellum or the hippocampus (Aggarwal et al., 2012; Lundell et al., 2015), as well as ischemia (Does et al., 2003; Aggarwal et al., 2014; Wu et al., 2018) and tumors (Colvin et al., 2008; Colvin et al., 2011; Xu et al., 2012; Reynaud et al., 2016). The gradient hardware of clinical MRI systems typically limits the frequency range to 50 Hz which remains sufficient to obtain useful contrast in human brains (Baron and Beaulieu, 2014; Van et al., 2014; Baron et al., 2015; Arbabi et al., 2020; Tetreault et al., 2020). The accessible frequency range is continuously being extended by further developments of gradient hardware (Tan et al., 2020; Hennel et al., 2021; Michael et al., 2022; Dai et al., 2023).

Interpretation of time- or frequency-dependent ADCs in terms of geometric properties is confounded by heterogeneity of the investigated object on length-scales larger than those being mixed by diffusional exchange during the motion-encoding gradients, for instance, multiple tissue types or WM fiber bundles with different orientations within the same imaging voxel. Gradient waveforms with successive diffusion encoding in multiple directions reduce or even remove the effects of diffusion anisotropy on the acquired signal (Mori and van Zijl, 1995) and, when combined with data from conventional unidirectional encoding, enable separation between isotropic and anisotropic sources of intravoxel heterogeneity (Eriksson et al., 2013; Lasič et al., 2014; Szczepankiewicz et al., 2015; Topgaard, 2016b; Westin et al., 2016). By capitalizing on the gain in information content obtained by varying the “shape” (Westin et al., 2014) of the *b*-tensor, the parametric diffusion tensor distribution (DTD) approach (Basser and Pajevic, 2003; Jian et al., 2007; Leow et al., 2009; Magdoom et al., 2021) for describing multi-component diffusion in anisotropic media has been generalized to nonparametric distributions (de Almeida Martins and Topgaard, 2016; Topgaard, 2019a) with in vivo applications in preclinical (Yon et al., 2020) and clinical MRI (Reymbaut et al., 2020b; Daimiel Naranjo et al., 2021). Incorporating the sensitivity to restriction of OGSE into the DTD framework (Lundell et al., 2019) yields nonparametric frequency-dependent DTDs or “**D**(*ω*)-distributions” (Narvaez et al., 2021).

Combining DTD with diffusion-relaxation correlation (Galvosas and Callaghan, 2010; Bernin and Topgaard, 2013; Song et al., 2016) via variable repetition- and/or echo times yields multidimensional correlations between **D** and the relaxation rates *R*_1_ and *R*_2_ (de Almeida Martins and Topgaard, 2018), which has been demonstrated in vivo on small-animal (Rosenberg et al., 2022) and whole-body MRI systems (de Almeida Martins et al., 2020; de Almeida Martins et al., 2021; Martin et al., 2021; Reymbaut et al., 2021). OGSE, tensor-valued encoding, and diffusion-relaxation correlation were recently combined into the overarching “massively multidimensional diffusion-relaxation correlation MRI” framework, yielding nonparametric **D**(*ω*)-*R*_1_-*R*_2_ distributions by Monte Carlo data inversion (Narvaez et al., 2022). While the high-performance gradient hardware of microimaging systems allows comprehensive exploration of both the spectral and tensorial aspects of **b**(*ω*) via “double rotation” gradients waveforms (Jiang et al., 2023), the modest gradient amplitude offered by clinical scanners limits the accessible frequencies to ranges that may not be sufficient to quantify the effects of restriction. Conversely, attempting frequency-independent inversion of data that features effects of restriction may result in systematic errors of the estimated parameters (de Swiet and Mitra, 1996; Jespersen et al., 2019).

In this article, we investigate fit residuals and bias of metrics extracted from nonparametric **D**(*ω*)-*R*_1_-*R*_2_ and **D**-*R*_1_-*R*_2_ distributions obtained with (Narvaez et al., 2022) and without (de Almeida Martins and Topgaard, 2018), respectively, inclusion of frequency-dependence in the inversion of experimental data covering wide and narrow frequency ranges. More specifically, we analyze data acquired as a function of echo time, repetition time, and diffusion-encoding gradient waveforms giving *b*-tensor anisotropies (Eriksson et al., 2015) *b*_Δ_ = –0.5, 0, and 1 using a microimaging system allowing a wide frequency range (35 to 320 Hz at *b* = 4·10^9^ sm^−2^) for phantoms with well-defined diffusion properties, excised tumor tissue, and fixated ex vivo mouse brain, as well as a conventional clinical system limited to a narrow frequency range (5 to 11 Hz at *b* = 3·10^9^ sm^−2^) for in vivo human brain. Building up for interpretation of the results on frequency-dependence of the water populations in the latter data, we go through the results for the series of simpler systems where mechanisms contributing to the mixing of water populations and frequency dispersion are well-known from the chemistry literature, an important message being that the gradient waveform durations and frequency ranges selected mainly because of hardware constraints most likely coincide with some processes in the continuous range of exchange and restriction mechanisms in the living human brain. Consequently, we suggest that complete or partial mixing of water populations should be considered when interpreting the obtained results and frequency-dependence should explicitly be included in the data inversion to accommodate the cases where restriction mechanisms happen to fall within the used frequency window.

## Methods

### Samples

Saturated salt solutions were prepared by adding Mg(NO_3_)_2_·6 H_2_O and Co(NO_3_)_2_·6 H_2_O (both from Sigma-Aldrich Sweden AB) to H_2_O (Milli-Q quality) until reaching the solubility limits 71 g Mg(NO_3_)_2_ and 97 g Co(NO_3_)_2_ per 100 mL H_2_O (Rumble, 2021). A small amount of Co(NO_3_)_2_ solution (0.27 wt%) was added to the Mg(NO_3_)_2_ solution to increase ^1^H_2_O *R*_1_ and *R*_2_ to approximately 2 and 20 s^−1^, respectively. A lamellar liquid crystal (Ekwall et al., 1969; Jiang et al., 2021) was prepared by mixing 85.79 wt% H_2_O (Milli-Q), 9.17 wt% 1-decanol (Sigma-Aldrich Sweden AB), and 5.04 wt% sodium octanoate (J&K Scientific via Th. Geyer in Sweden). The composite phantom was assembled by inserting 4-mm NMR tubes containing salt solution and liquid crystal into a 10-mm NMR tube with H_2_O.

The excised tumor tissue was obtained by culture of human neuroblastoma cells grown at 37 °C and 5% CO_2_ in a complete medium (RPMI 1640 supplemented with 10% fetal bovine serum and 1% penicillin/streptomycin). Approximately 2·10^6^ of those tumor cells were subcutaneously inoculated to a female BALB/c mouse (Janvier Labs, France). The mouse was sacrificed after 5 weeks of tumor growth and the tumor was removed and immediately transferred to a 10-mm NMR tube containing 4% PFA in phosphate buffer solution (Histolab, Sweden). The sample was stored at room temperature for several years before being investigated with MRI.

The ex vivo mouse brain was obtained from an 8-week-old C57BL/6 female mouse intracardially perfused with 0.9% saline solution followed by 4% paraformaldehyde (PFA) fixation. The brain was carefully extracted from the skull and stored in 2% PFA solution at 4°C before MRI acquisition. The procedure was approved by the Animal Committee of the Provincial Government of Southern Finland following the guidelines established by the European Union Directives 2010/63/EU.

The human data were obtained on a healthy young adult with approval from the local institutional review board and informed consent of the patient.

### MRI acquisition and reconstruction

The composite phantom, excised tumor tissue, and ex vivo mouse brain were investigated using a Bruker Avance Neo spectrometer (Bruker Biospin, Karlsruhe, Germany) with an 11.7 T magnet, a MIC-5 probe delivering 3 Tm^−1^ maximum gradient strength, and a 10 mm ^1^H radiofrequency coil. Images were acquired in Paravision 360 v1.1 with custom-made sequences based on either Rapid Acquisition with Relaxation Enhancement (RARE) (Hennig et al., 1986) or multi-slice multi-echo (MSME) (Edelstein et al., 1980) with spin-echo diffusion preparation according to the generic pulse sequence scheme presented in Figure 1a. The mouse brain MSME images were acquired at 25 °C sample temperature with 14×9×0.5 mm^3^ field of view (FOV) and 100×64×1 reconstructed image size, giving 140×140×500 µm^3^ resolution. Using a single scan per each of the 36 phase encoding steps with 1.8 partial Fourier factor led to an acquisition time of 36 hours and 8 minutes. The phantoms and tumor RARE images were acquired at 20 °C sample temperature with 12×12×0.5 mm^3^ FOV and 64×64×1 matrix size, giving 190×190×500 µm^3^ resolution. A partial Fourier factor of 1.8 was used in the first phase dimension to reduce the echo time and allow single-shot acquisition with 36 echoes. Using four averages led to an acquisition time of 4 hours and 7 minutes. The images were reconstructed with Paravision 360 v1.1, followed by denoising (Cordero-Grande et al., 2019) implemented in MRTrix3 (Tournier et al., 2019), as well as Gibbs ringing removal (Kellner et al., 2016) for the mouse brain dataset.

**Figure 1:**
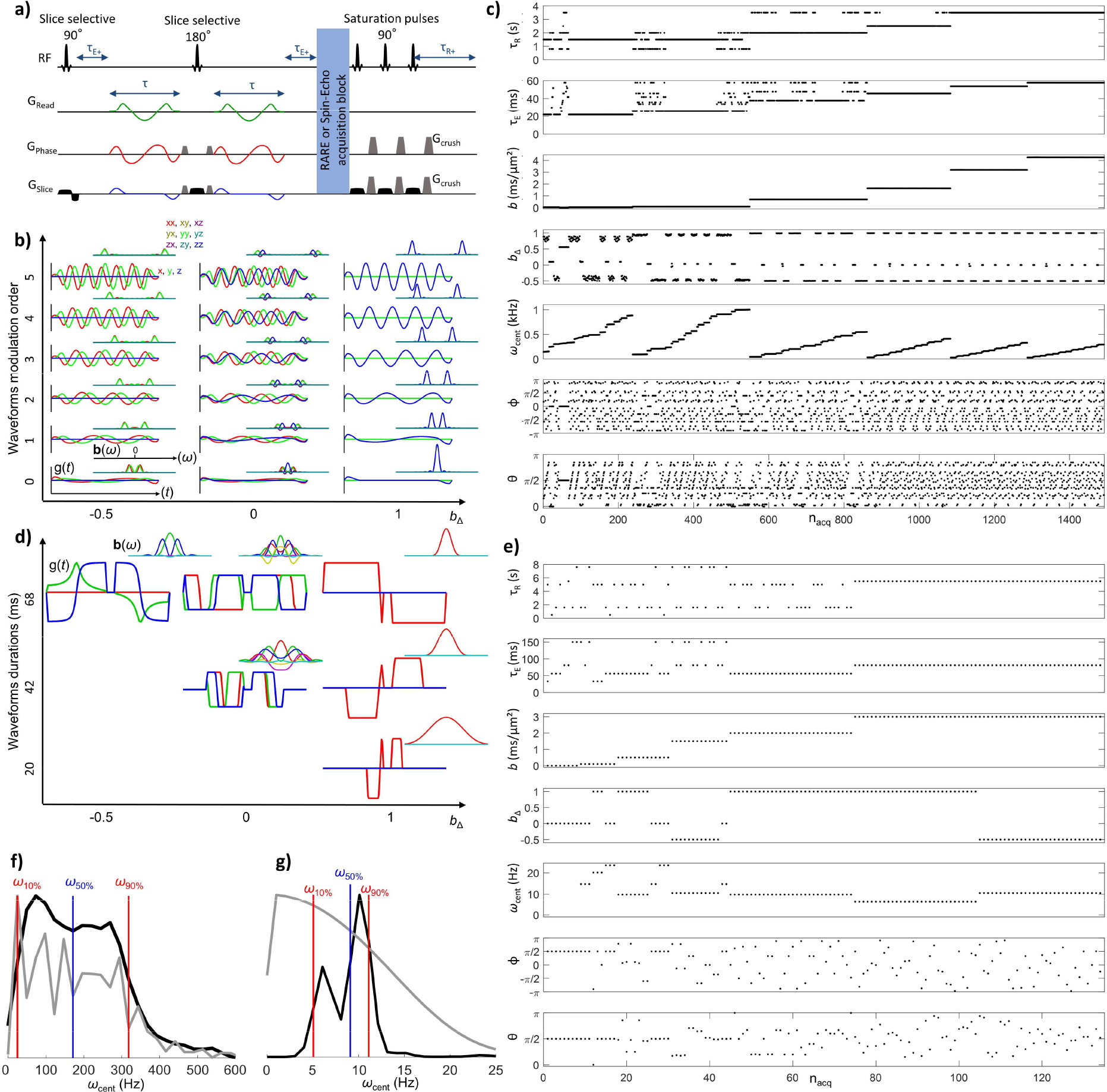
a) Generic pulse sequence scheme for multidimensional diffusion-relaxation correlation MRI integrating variable echo time *τ*_E_, recovery time *τ*_R_, and time-modulated gradient waveforms **g**(*t*). b) Double-rotation waveforms and encoding spectra **b**(*ω*), calculated with Eqs. (1)-(3), on the top right used for the preclinical acquisitions for encoding anisotropy *b*_Δ_ = –0.5, 0, and 1, defined in Eq. (8), and up to five oscillations. c) Preclinical (wide *ω*_cent_-range) acquisition protocol with 1491 images labeled with acquisition number *n*_acq_ and sorted by *b*-value and centroid frequency *ω*_cent_, defined in Eqs. (6) and (7), respectively. d) Numerically optimized gradient waveforms used for clinical acquisition. e) Clinical (narrow *ω*_cent_-range) acquisition protocol with 134 image volumes. f) Total spectral content of the preclinical protocol (gray line) and *b*-weighted *ω*_cent_-distribution (black line). Red and blue vertical lines indicate the 10th, 50th, and 90th percentiles of the *ω*_cent_-distribution: *ω*_10%_ = 35 Hz, *ω*_50%_ = 190 Hz, and *ω*_90%_ = 320 Hz. g) Total spectral content and *ω*_cent_-distribution of the clinical protocol shown as in panel f, yielding *ω*_10%_ = 5 Hz, *ω*_50%_ = 9 Hz, and *ω*_90%_ = 11 Hz.

The in vivo data was recorded using a Siemens Magnetom Prisma (Siemens Healthineers AG, Erlangen, Germany) with a 3 T magnet, a gradient system providing a maximum amplitude of 0.08 Tm^−1^, and a 20-channel head coil. The images were acquired with a single-shot spin echo-echo planar imaging (SE-EPI) sequence customized for general gradient waveforms (Wetscherek et al., 2015; Martin et al., 2020). The acquisition parameters were: 230×230 mm^2^ FOV, 3 mm^3^ isotropic resolution, 30 slices in axial orientation, 1496 Hz/Px readout bandwidth, and a factor 3 acceleration with GRAPPA reconstruction. The total measurement time was 20 minutes. The data were preprocessed with denoising (Cordero-Grande et al., 2019; Tournier et al., 2019), removal of Rician noise baseline (Koay and Basser, 2006) and Gibbs ringing (Kellner et al., 2016), and motion and eddy current correction (Klein et al., 2010; Nilsson et al., 2015).

### Multidimensional diffusion and relaxation encoding

As illustrated in Figure 1, data were recorded for numerous combinations of recovery time *τ*_R_ and echo time *τ*_E_, encoding for longitudinal and transverse relaxation, as well as gradient waveforms **g**(*t*) targeting the frequency-dependence and anisotropy of the translational motion. The diffusion encoding properties are captured in the tensor-valued encoding spectrum **b**(*ω*) given by **g**(*t*) via the time-dependent dephasing vector **q**(*t*) and its Fourier transform **q**(*ω*) according to (Lundell and Lasič, 2020):

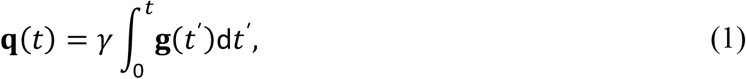

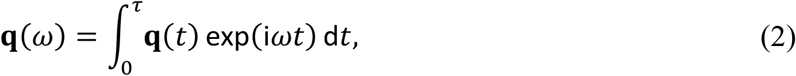

and

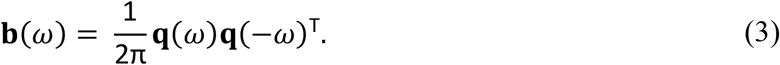

In the equations above, *γ* is the gyromagnetic ratio, *τ* is the of duration of the gradient waveform including the imaging gradients, and T denotes a matrix transpose. The conventional encoding power spectrum *b*(*ω*) (Callaghan and Stepišnik, 1995), encoding tensor **b** (Basser et al., 1994), and *b*-value (Le Bihan et al., 1986) are obtained from **b**(*ω*) via

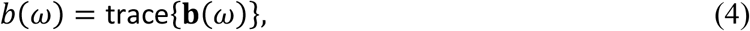

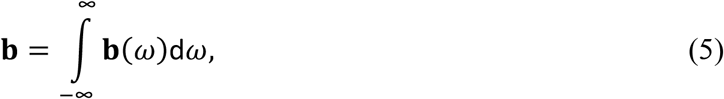

and

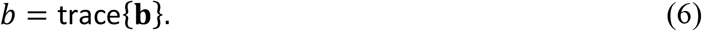

Although all spectral and tensorial aspects of **b**(*ω*) are used in the data inversion described below, for bookkeeping it is convenient to summarize the sensitivity to restriction and anisotropy in terms of the centroid frequency *ω*_cent_ (Ligneul and Valette, 2017; Arbabi et al., 2020) and anisotropy *b*_Δ_ (Eriksson et al., 2015), respectively, defined as

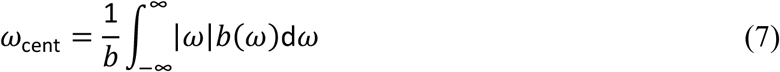

and

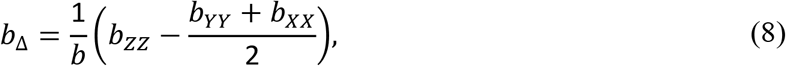

where *b*_*XX*_, *b*_*YY*_, and *b*_*ZZ*_ are the eigenvalues of **b** ordered according to the convention |*b*_*ZZ*_ – *b*/3| > |*b*_*XX*_ – *b*/3| > |*b*_*YY*_ – *b*/3| (Topgaard, 2016b, 2017). The directionality of the encoding is reported as the polar and azimuthal angles, Θ and Φ, of the eigenvector corresponding to the *b*_*ZZ*_ eigenvalue.

In the preclinical system, diffusion encoding was performed with two identical self-refocusing gradient waveforms, while in the clinical setting, the waveforms were different and not self-refocused to maximize the *b*-value per encoding time unit (Sjölund et al., 2015). Gradient waveforms and the corresponding encoding spectra **b**(*ω*) at *b*_Δ_ = –0.5, 0, and 1 are shown in Figure 1b and d for the preclinical and clinical scanners, respectively. In the preclinical setting, the gradient waveforms were generated by double rotation of the *q*-vector (Jiang et al., 2023) and the *ω*_cent_ dimension was explored over a wide range by varying the waveform duration from 4 to 22 ms and the number of oscillations from 0 to 5 as shown in Figure 1b. The values of *b* and *ω*_cent_ for each acquired image volume are shown in Figure 1c, illustrating that the highest values of *ω*_cent_ are achieved for low *b*-values only. The frequency-modulated gradient waveforms were normalized to give constant *b*-values at identical waveform lengths allowing us to map the entire *b*-value space (from 0.033 to 4.25·10^9^ sm^−2^) with all the waveforms. The deviations of *b*_Δ_ from the target values –0.5, 0, and 1 at low *b*-values result from the imaging gradients which were all taken into account when computing **b**(*ω*) via Eqs. (1)-(3) above. In the clinical setting, the limited maximum gradient amplitude necessitated numerical optimization of the waveforms (Sjölund et al., 2015) to reach *b* = 3·10^9^ sm^−2^ at *τ*_E_ = 83 ms and *b*_Δ_ = –0.5, 0, and 1 as previously used in Refs. (Martin et al., 2020) and (Martin et al., 2021). The directional dependence of the diffusion was probed by rotating the waveforms in Figure 1b and d according to the angles Θ and Φ in Figure 1c and e.

Figure 1f and g show quantitative assessments of the range of frequencies investigated in each acquisition protocol in terms of the *b*-weighted *ω*_cent_-distribution and the total spectral content expressed as the sum of *b*(*ω*) over all acquisitions with index *n*_acq_. For the preclinical protocol (Figure 1f), each waveform yields *b*(*ω*) focused on a narrow frequency range centered on *ω*_cent_ and the width of the total spectral content matches the one of the broad *ω*_cent_-distribution, implying that the protocol is appropriate for investigating frequency-dependence. Conversely, each clinical waveform yields a poorly defined *b*(*ω*) with width that surpasses the spread of *ω*_cent_ across the acquisitions, resulting in total spectral content far broader than the narrow *ω*_cent_-distribution (Figure 1g) and data that are less amenable for extracting frequency-dependence.

The preclinical protocol of 1491 images includes variable *τ*_R_ between 0.8 and 3.5 s as well as variable *τ*_E_ between 13 and 49 ms for MSME and 22 to 58 ms for RARE, the latter being displayed in Figure 1c. The clinical protocol with 134 image volumes includes *τ*_R_ from 0.5 to 7.6 seconds and *τ*_E_ from 33 and 150 ms. Values of *τ*_R_ below 1.6 s were reached by acquiring the 30 slices in packages with a few slices each.

### Monte Carlo data inversion

The diffusion- and relaxation encoded signal *S*[**b**(*ω*),*τ*_R_,*τ*_E_] is expressed as the sum of components *i* characterized by their weights *w*_i_, tensor-valued diffusion spectra **D**_*i*_(*ω*), and relaxation rates *R*_1,*i*_ and *R*_2,*i*_ according to (Narvaez et al., 2022)

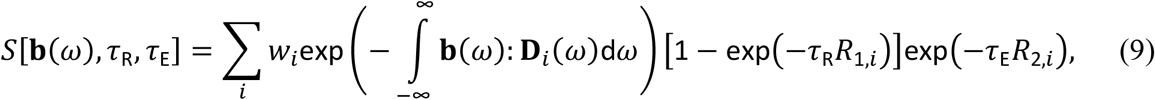

where the colon indicates a generalized scalar product (Kingsley, 2006). In the special case of constant **D**_*i*_(*ω*) = **D**_*i*_ in the investigated frequency interval, Eq. (9) reduces to (de Almeida Martins and Topgaard, 2018)

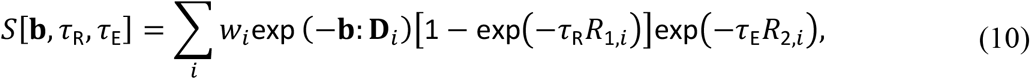

where the relation between **b**(*ω*) and **b** is given in Eq. (5). We emphasize that the variables *b, ω*_cent_, *b*_Δ_, Θ, and Φ—familiar from the literature and reported in the acquisition protocols in Figure 1—are here used only for bookkeeping and are not explicitly included in the expressions Eqs. (9) and (10) applied in the data inversion.

For computational convenience, the component tensors are assumed to have axial symmetry:

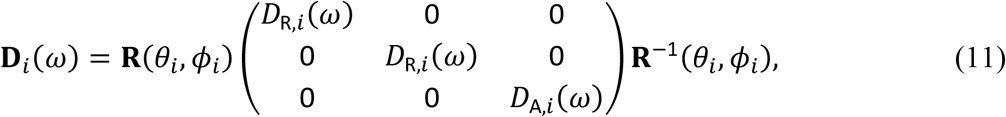

where **R**(*θ*,*ϕ*) is a rotation matrix. Further, the axial and radial eigenvalues *D*_A,*i*_(*ω*) and *D*_R,*i*_(*ω*) are approximated as Lorentzians (Narvaez et al., 2021),

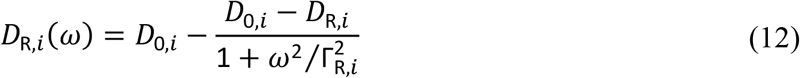

and

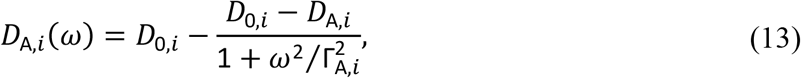

where *D*_A,*i*_ and *D*_R,*i*_ are the low-*ω* diffusivities in the axial and radial directions, respectively, *D*_0,i_ is the high-*ω* diffusivity, assumed to be isotropic, and Γ_A,*i*_ and Γ_R,*i*_ are the values of *ω* at the mid-points of the transitions. The *ω*-independent case in Eq. (10) corresponds to *D*_A,*i*_(*ω*) = *D*_A,*i*_ and *D*_R,*i*_(*ω*) = *D*_R,*i*_.

A Monte Carlo algorithm implemented in the *md-dmri* (Nilsson et al., 2018) Matlab toolbox was used to estimate ensembles of discrete distributions in the spaces [*D*_A_,*D*_R_,*θ,ϕ,D*_0_,Γ_A_,Γ_R_,*R*_1_,*R*_2_] (Narvaez et al., 2022) and [*D*_A_,*D*_R_,*θ,ϕ,R*_1_,*R*_2_] (de Almeida Martins and Topgaard, 2018), corresponding to Eqs. (9) and (10), respectively. Using the terminology in Reymbaut et al. (Reymbaut et al., 2020a), the inversion was performed with 20 steps of proliferation, 20 steps of mutation/extinction, 200 input components per step of proliferation and mutation/extinction, and 10 output components. Bootstrapping was performed by 100 repetitions using random sampling with replacement. Preclinical (clinical) data was inverted with the parameter limits 5·10^−12^ m^2^s^−1^ < *D*_0/A/R_ < 5·10^−9^ m^2^s^−1^, 0.1 s^−1^ < Γ_A/R_ < 10^5^ s^−1^, 0.1 s^−1^ < *R*_1_ < 4 s^−1^, and 4 s^−1^ < *R*_2_ < 150 s^−1^ (5·10^−11^ m^2^s^−1^ < *D*_0/A/R_ < 5·10^−9^ m^2^s^−1^, 0.1 s^−1^ < Γ_A/R_ < 10^4^ s^−1^, 0.2 s^−1^ < *R*_1_ < 2 s^−1^, and 1 s^−1^ < *R*_2_ < 30 s^−1^).

### Quantitative parameter distributions and maps

To enable visualization, the distributions in the primary analysis space [*D*_A_,*D*_R_,*θ,ϕ,D*_0_,Γ_A_,Γ_R_,*R*_1_,*R*_2_] were evaluated at selected values of *ω*, using Eqs. (12) and (13), and projected onto the dimensions of isotropic diffusivity *D*_iso_(*ω*) and squared normalized diffusion anisotropy *D*_Δ_^2^(*ω*) (Conturo et al., 1996; Eriksson et al., 2015; Topgaard, 2019a) via

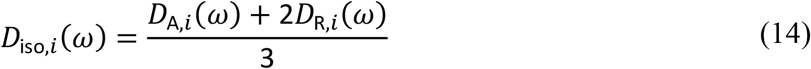

and

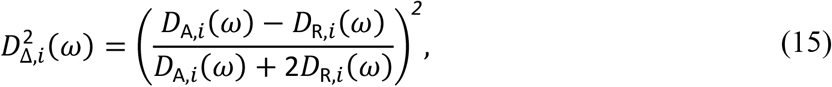

respectively, yielding *ω*-dependent distributions in the [*D*_iso_(*ω*),*D*_Δ_^2^(*ω*),*θ,ϕ,R*_1_,*R*_2_] space. Projections onto the 2D *D*_iso_-*D*_Δ_^2^ plane (Topgaard, 2019a) were obtained by mapping the weights *w*_*i*_ of the discrete components onto a 64×64 mesh using a 3×3 grid points Gaussian kernel. Image segmentation was performed by dividing the 2D *D*_iso_-*D*_Δ_^2^ space into three bins with diffusion properties characteristic for white matter (WM, bin1), grey matter (GM, bin2), and cerebrospinal fluid (CSF, bin 3) using the limits bin1: *D*_iso_ < 1·10^−9^ m^2^s^−1^ and *D*_Δ_^2^ > 0.25; bin2: *D*_iso_ < 1·10^−9^ m^2^s^−1^ and *D*_Δ_^2^ < 0.25; and bin3: *D*_iso_ > 1·10^−9^ m^2^s^−1^ (bin1: *D*_iso_ < 2·10^−9^ m^2^s^−1^ and *D*_Δ_^2^ > 0.25; bin2: *D*_iso_ < 2·10^−9^ m^2^s^−1^ and *D*_Δ_^2^ < 0.25; and bin3: *D*_iso_ > 2·10^−9^ m^2^s^−1^) for the ex vivo mouse (in vivo human) data and calculating bin-resolved signal fractions *f*_bin*n*_ by

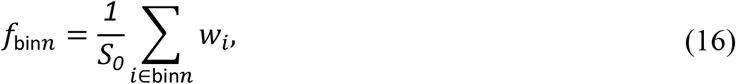

where

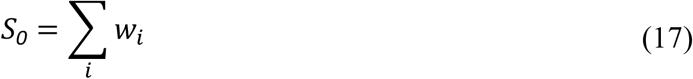

is the signal extrapolated to *b* = 0, *τ*_R_ = ∞, and *τ*_E_ = 0. The signal fractions were converted to RGB color via

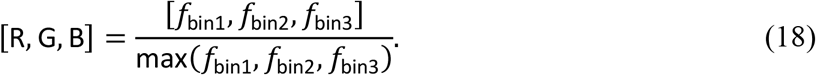

The rich information in the [*D*_iso_(*ω*),*D*_Δ_^2^(*ω*),*θ,ϕ,R*_1_,*R*_2_]-distributions was further condensed into means E[*X*] over selected dimensions according to (Topgaard, 2019a)

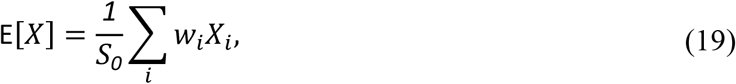

where *X* symbolizes *D*_iso_(*ω*) and *D*_Δ_^2^(*ω*) at the frequencies *ω*_10%_, *ω*_50%_, and *ω*_90%_ indicated in Figure 1f and g, as well as *R*_1_ and *R*_2_. The dispersion of the diffusion metrics within the investigated frequency window Δ_*ω*/2π_E[*X*] was calculated through (Aggarwal et al., 2012; Narvaez et al., 2021)

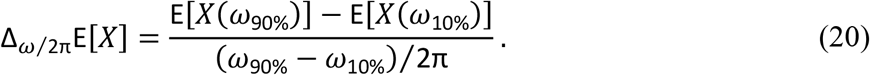

To evaluate the uncertainty of the data inversion procedure, the diffusion- and relaxation encoded signals *S*[**b**(*ω*),*τ*_R_,*τ*_E_], 2D *D*_iso_-*D*_Δ_^2^ projections, extrapolated signals *S*_0_, signal fractions *f*_bin*n*_, means E[*X*], and dispersions Δ_*ω*/2π_E[*X*] were calculated independently for each of the 100 bootstrap replicates. The values underlying the graphs and maps in the following figures were obtained as medians over these 100 replicates.

For the *ω*-independent analysis based on Eq. (10), the metrics in Eqs. (14)-(19), including the binning, was performed with the *ω*-independent values of *D*_A_ and *D*_R_ obtained directly in the primary analysis space [*D*_A_,*D*_R_,*θ,ϕ,R*_1_,*R*_2_]. Comparison between the *ω*-dependent and *ω*-independent results was performed by evaluating the former at the center of the investigated frequency window, corresponding to the value *ω*_50%_ labeled in Figure 1f and g, and computing the normalized difference via

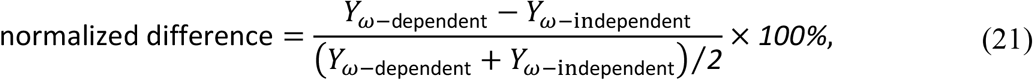

where *Y* represents *S*_0_, E[*D*_iso_], E[*D*_Δ_^2^], E[*R*_1_], E[*R*_2_], or *f*_bin*n*_.

## Results and Discussion

Figure 2 shows data for a series of regions of interest (ROIs) in samples with distinct restriction and anisotropy characteristics investigated using the preclinical protocol shown in Figure 1c. The selected samples represent chemically “simple” systems, where the relevant time-scales and mechanisms affecting the observed diffusivities are well understood from the chemistry literature, as well as a series of biomedically more interesting tissue ROIs where the contributing mechanisms are the same as for the previous examples, but the relevant time scales are more difficult to predict due to the increased chemical complexity and the continuous range of structural organization levels from the molecular to the macroscopic.

**Figure 2:**
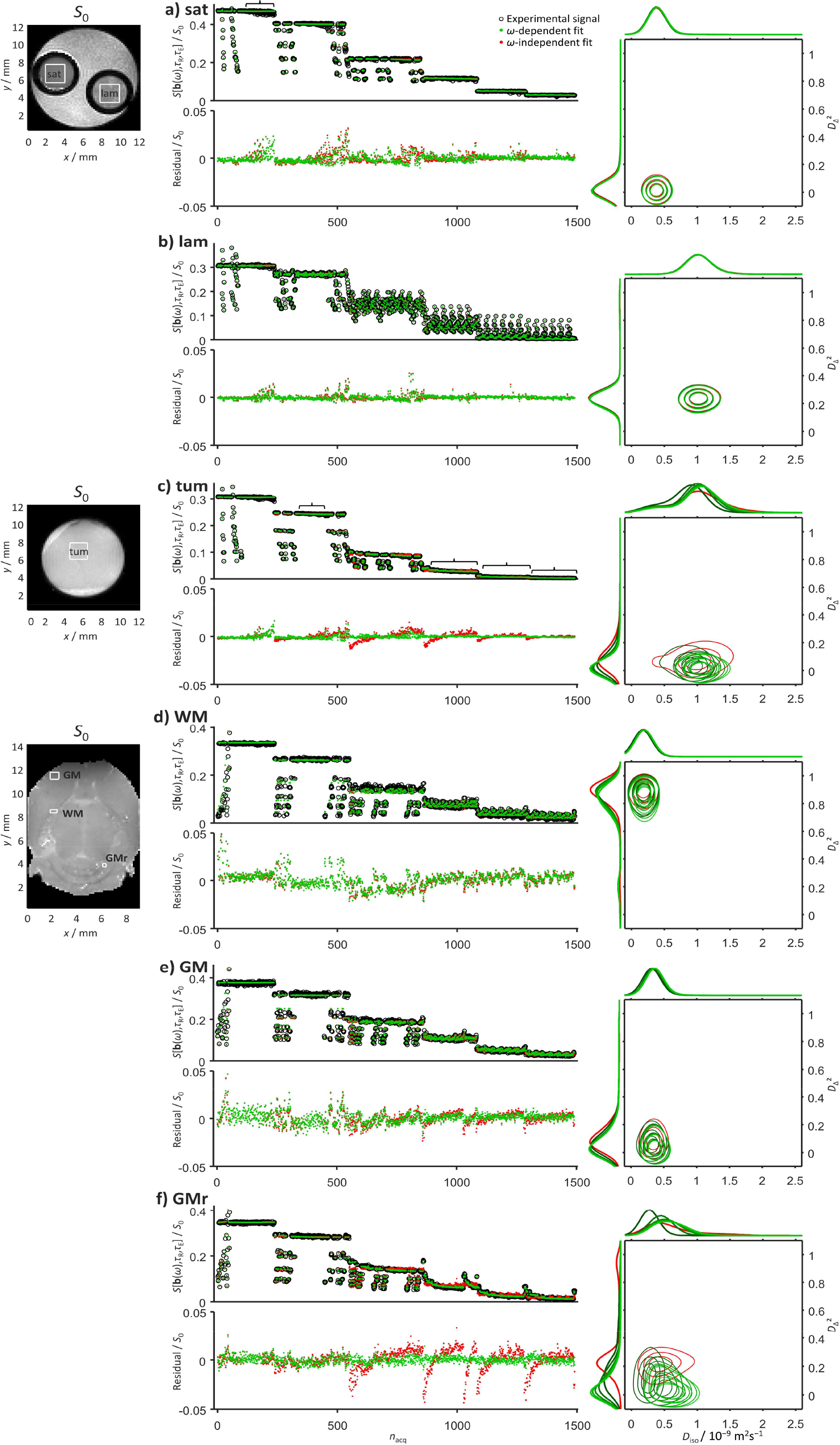
Comparison between *ω*-dependent and *ω*-independent data inversion results for illustrative cases with distinct water diffusion properties. a) Isotropic Gaussian diffusion in an aqueous solution saturated with magnesium nitrate salt (sat). b) Planar anisotropic Gaussian diffusion in a lamellar liquid crystal (lam). c) Isotropic restricted diffusion in tumor tissue (tum). d) Linear anisotropic Gaussian diffusion in white matter (WM) of the internal capsule. e) Isotropic restricted diffusion in the gray matter (GM) of the cortex. f) Isotropic restricted diffusion in the gray matter of the cerebellum (GMr). Panels to the left show labeled regions of interest (ROIs) for the composite phantom (lam and sat), excised tumor (tum), and ex vivo mouse brain (WM, GM, and GMr) on maps of the signal *S*_0_ extrapolated to *b* = 0, *τ*_R_ = ∞, and *τ*_E_ = 0, see Eq. (17). The center panels display measured signals *S*[**b**(*ω*),*τ*_R_,*τ*_E_] vs. acquisition number *n*_acq_ according to the preclinical (wide *ω*_cent_-range) protocol in Figure 1c (black circles), signals back-calculated from the distributions obtained by Monte Carlo inversion of the *ω*-dependent (green dots) and *ω*-independent (red dots) expressions in Eqs. (9) and (10), respectively, as well as residuals given by the differences between the measured and back-calculated signals. Signals and residuals are normalized with *S*_0_. The right part of the figure presents the *ω*-dependent (green) and *ω*-independent (red) distributions as projections onto the 2D *D*_iso_-*D*_Δ_^2^ plane (contour plots) as well as the 1D *D*_iso_ and *D*_Δ_^2^ dimensions (horizontal and vertical line plots sharing axes with the 2D plots). The *ω*-dependence is illustrated by overlaying color-coded plots for 5 linearly spaced values of *ω* between 35 (dark green) and 320 Hz (pale green). Overbraces in panels a and c point out protocol sections at constant *b, τ*_R_, and *τ*_E_ where the residuals are unaffected or decrease, respectively, by including *ω*-dependence in the inversion.

According to the literature, the saturated magnesium nitrate solution in panel a exhibits a water diffusivity of 0.44·10^−9^ m^2^s^−1^ at 25 ºC (Wadsö et al., 2009), which is 20% of the value 2.3·10^−9^ m^2^s^−1^ for pure water (Mills, 1973) on account of interactions between the water and the ions. The chemical composition reported in the methods section can be converted to a molar ratio of twelve water molecules per magnesium ion and two nitrate ions, implying that every water molecule is in direct atomic-level contact with neighboring ions. The water-ion electrostatic interactions are dominated by the magnesium ion because of its higher charge density, two positive charges for an ionic radius of 86 ppm, compared to the nitrate ion with one negative charge for a thermochemical radius of 179 ppm (Simoes et al., 2017). In dilute solution, the first hydration shell of the magnesium ion comprises six water molecules with a lifetime of about 1 μs before exchange with the less well-defined and more labile second hydration shell and surrounding bulk water (Neely and Connick, 1970; Bleuzen et al., 1997). Including the < 10^−12^ s decay of the velocity autocorrelation function for pure water (Balucani et al., 1996) and the > 10^3^ s time required for the mean squared displacement to equal the distance between the walls of the 4 mm glass tube, we would thus expect the diffusion spectrum **D**(*ω*) to show *ω*-dependence at the widely space values 10^−3^, 10^6^, and 10^12^ Hz, but not within the ~30-300 Hz range defined by the currently used gradient waveforms with ~50 ms total duration. Returning to Figure 2a, the expectations of isotropic Gaussian diffusion are borne out by the absence of signal modulations from the acquisition variables *ω*_cent_, *b*_Δ_, Θ, and Φ at constant *b, τ*_R_, and *τ*_E_, as well as nearly identical fit residuals for data inversions based on the *ω*-dependent and *ω*-independent expressions Eqs. (9) and (10), respectively. The corresponding 2D *D*_iso_-*D*_Δ_^2^ projections of the obtained distributions comprise a single peak at *D*_iso_ = 0.4·10^−9^ m^2^s^−1^ and *D*_Δ_^2^ = 0 with no detectable *ω*-dependence. The peak width originates mainly from the variability of the 100 replicate solutions obtained by the bootstrapping and Monte Carlo data inversion (Reymbaut et al., 2020a), which includes neither the conventional Tikhonov regularization (Provencher, 1982; Whittal and MacKay, 1989), leading to peak broadening, nor sparsity constraints (Berman et al., 2013; Aranda et al., 2015) favoring narrow peaks. Our results at 20 ºC are consistent with literature data at 25 ºC (Wadsö et al., 2009) and should be interpreted as an average over exchanging water populations in the first and second hydration shells of the magnesium ions with no influence from restriction by the glass tube walls. More detailed comparison between the plots of residuals in Figure 2 and the protocol in Figure 1c shows that acquisitions above 500 Hz yield data that cannot be fully captured even by the *ω*-dependent expression in Eq. (9) despite the *b*-value being too low to give any appreciable diffusion weighting, indicating that these data points are corrupted by image artifacts with magnitude of a few percent that may be difficult to detect by visual inspection of the raw images. Although these data points (overbrace in panel a) should be excluded in improved versions of the protocol, they appear to have negligible influence on the obtained distributions and are useful in the context of this work as a reference for whether or not inclusion of *ω*-dependence in the inversion improves the analysis.

The lamellar liquid crystal in Figure 2b consists of stacks of decanol and sodium octanoate bilayers separated by water. The bilayers have a thickness of 2.5 nm (Ekwall et al., 1969), a lamellar repeat distance of 15 nm (Jiang et al., 2021), and are organized with the hydrophilic hydroxyl and carboxylate groups facing the water and hydrophobic hydrocarbon chains in the bilayer interior. The sodium ions are distributed across the 12.5 nm thick water layers with preferential location within a few nm distance from the bilayer surface on account of electrostatic interactions with the oppositely charged carboxylate groups (Evans and Wennerström, 1999). There are six water molecules within the first hydration shells of both divalent magnesium and monovalent sodium ions, but the lifetime is only ~1 ns (Helm and Merbach, 1999) for the latter species because of the lower charge density (one positive charge for an ionic radius of 116 pm). Additionally, proton exchange between decanol hydroxyl groups and water takes place on time scales lower than 1 ms (Hills, 1990). The chemical composition of the liquid crystal corresponds to ~160 water and two decanol molecules per sodium and octanoate ion pair, and, as opposed to the case of the saturated salt solution, only a small fraction of the water molecules is in direct atomic-scale contact with the ions or bilayers. The low water concentration within the hydrophobic interior of the bilayers makes them efficient barriers for water diffusion (Evans and Wennerström, 1999), and the time for the mean squared displacement to cover the gap between two adjacent bilayers is ~40 ns. The lateral extension of the bilayers may approach macroscopic length scales and is ultimately limited by the walls of the glass tube (Bernin et al., 2014; Topgaard, 2016a), leading to characteristic time scales above 10^2^ s for diffusional exchange between differently oriented bilayer sections having a radius of curvature above 1 mm (Lutti and Callaghan, 2007). Partial alignment of the chain-like decanol molecules and octanoate ions in the direction of the bilayer normal vector renders the motional averaging of intermolecular ^1^H-^1^H dipolar couplings incomplete (Wennerström, 1973), leading to transverse relaxation on time scales shorter than the ~20 ms minimum echo time in the current protocol and minimal contribution from these species to the intensity of the detected images. Taken together, we may thus anticipate *ω*-dependence of **D**(*ω*) at multiple frequencies including 10^−2^ (water diffusion along bilayer curvature), 10^3^ (water-decanol chemical exchange), 10^8^ (diffusion across water layers), 10^9^ (lifetime of water in sodium ion hydration layer), and 10^12^ Hz (transition from ballistic to diffusive regime of pure water), none of which being located within the narrow *ω*-range explored with the present gradient waveforms. Expectedly, the data for the liquid crystal in Figure 2b shows pronounced signal modulations as a function of the *b*_Δ_, Θ, and Φ acquisition variables, which by itself indicates anisotropy, but no differences in fit residuals between the *ω*-dependent and *ω*-independent inversions, showing that the diffusion is Gaussian in the investigated window. The highest fit residuals of a few percent are found at low-*b* and high-*ω*_cent_ acquisitions and presumably originate from the subtle image artifacts previously discussed for the salt solution. The 2D *D*_iso_-*D*_Δ_^2^ projections feature a single *ω*-independent peak at *D*_iso_ = 1.0·10^−9^ m^2^s^−1^ and *D*_Δ_^2^ = 0.25, corresponding to *D*_A_ << *D*_R_ and *D*_R_ = 1.5·10^−9^ m^2^s^−1^. The latter value describes lateral diffusion along the planes of the bilayers and is given by a ~50 ms time average over protons in multiple exchanging populations including pure water in the center of the water layers, water in the hydration shells of the sodium ions and hydrophilic groups at the surfaces of the bilayers, and the hydroxyl groups of the decanol molecules. Additionally, the diffusion of the water molecules may be hindered by the roughness of the hydrophobic-hydrophilic interface originating from molecular protrusions (Ben-Shaul and Gelbart, 1985) or bilayer undulations (Lindahl and Edholm, 2000), leading to an observed lateral diffusivity that is 75% of the value 2.0·10^−9^ m^2^s^−1^ for pure water at 20 ºC (Holz et al., 2000).

The data in Figure 2c is obtained on excised tumor tissue preserved in paraformaldehyde solution. Despite the increase in chemical and structural complexity, the basic mechanisms contributing to lowering the observed diffusivity compared to the reference state of pure water are the same as for the simpler systems in panels a and b, namely proton exchange between water and labile functional groups, hydration of ions, and obstruction by larger molecules and aggregates. While the importance of the two first mechanisms could in principle be estimated by detailed analysis of the chemical composition of the tissue sample (Persson and Halle, 2008), the experimental observations are dominated by the latter mechanism which depends critically on the details of how the molecules are spatially arranged—in particular the assembly of lipids into bilayers that may or may not be efficient barriers for water diffusion (Topgaard, 2020). From the perspective of water dynamics, biological tissues can conceptually be divided into numerous subvolumes with different local concentrations of everything from small ions and metabolites to large macromolecules and macromolecular assemblies that influence water motion via the mechanisms of hydration and obstruction. Some of these subvolumes are formed by the thermodynamic equilibrium mechanism of liquid-liquid phase separation into regions with low and high concentration of macromolecules (Hyman et al., 2014; Banani et al., 2017). Other subvolumes are formed by semipermeable biomembranes that encapsulate regions of space with different chemical compositions than the surroundings and prevent equilibration of concentration gradients. The subvolumes—with or without biomembrane enclosure—have dimensions evenly spread out within the 10 nm to 100 μm range that is of relevance for rationalizing diffusion data, some examples being vesicles, condensate droplets, lysosomes, mitochondria (with internal compartmentation), endoplasmic reticulum, nucleus, cytosol, and the extracellular space. Because of the varying barrier properties of the biomembranes and the multiple structural levels in the tumor tissue, we may expect diffusional exchange of water between the subvolumes on a continuous range of time scales as well as a continuous *ω*-dependence of **D**(*ω*) filling in the gaps of the numerous processes from 10^−2^ to 10^12^ Hz described above for the salt solution and lamellar liquid crystal. In some aspects, the signal data for the tumor in panel c resembles the one from the salt solution in panel a, showing minor influence of the acquisition variables Θ and Φ at constant *b, τ*_R_, and *τ*_E_, consistent with isotropic diffusion, as well as elevated residuals originating from image artifacts in the low *b* and excessively high *ω*_cent_ range of the data. At higher *b*, the tumor data displays a marked dependence of the signal on *ω*_cent_ and greatly improved fit residuals with the *ω*-dependent analysis (overbrace in panel c) indicating *ω*-dependence of **D**(*ω*) within the investigated window ~30-300 Hz. The corresponding *ω*-dependent 2D *D*_iso_-*D*_Δ_^2^ projections show a single peak moving from E[*D*_iso_] = 0.84·10^−9^ m^2^s^−1^ and E[*D*_Δ_^2^] = 0.050 at 35 Hz to E[*D*_iso_] = 1.1·10^−9^ m^2^s^−1^ and E[*D*_Δ_^2^] = 0.014 at 320 Hz. Although the observed values for E[*D*_iso_] can be reproduced by inserting 1.2·10^−9^ m^2^s^−1^ local diffusivity and 7 μm radius in the model for restricted diffusion in a closed spherical compartment (Stepišnik, 1993), we emphasize that such a geometric interpretation is most certainly an oversimplification that may give rise to misconceptions if applied by users not familiar with the underlying assumptions and the plethora of alternative models with equal ability to describe the experimental results. Without overinterpretation, we may state that all detectable proton populations with potentially distinct diffusion properties are mixed on the ~50 ms time scale of the diffusion encoding gradients, giving rise to a single peak in the 2D *D*_iso_-*D*_Δ_^2^ projections, and that these exchange-averaged populations experience nearly isotropic structural barriers on the length scale of a few micrometers. Converting these imprecise statements to quantitative information about, e.g., compartment sizes and shapes, barrier permeabilities, and local diffusivities, would require model assumptions that are difficult to justify in light of the known chemical and structural complexity.

The remaining panels d, e, and f in Figure 2 present result for three regions of the ex vivo mouse brain: white matter (WM) in the internal capsule, gray matter in the cortex (GM), and gray matter in the cerebellum (GMr). WM consists of closely packed aligned axons with ~0.1-10 μm diameters (Saliani et al., 2017) and lengths on the macroscopic scale. The axons are piecewise enclosed by myelin sheaths formed by multilayer wraps of oligodendrocyte cell membranes. In addition to the numerous cellular-scale subvolumes, WM can on a coarser level be divided into the intraaxonal and extracellular spaces, as well as the myelin sheaths which in themselves have intra- and extracellular spaces. All these subvolumes have distinct types of organization of macromolecular assemblies and biomembranes that determine the local diffusion properties of the water. In analogy with the reasoning for the tumor tissue, we may expect some exchange or restricted diffusion process to occur at any given time scale. The data for WM in panel d shares some distinguishing features with the liquid crystal in Figure 2b: signal fluctuations with acquisition variables *b*_Δ_, Θ, and Φ at constant *b, τ*_R_, and *τ*_E_, but only minor differences in fit residuals between the *ω*-dependent and *ω*-independent inversions. These observations indicate anisotropic diffusion without detectable *ω*-dependence within the investigated ~30-300 Hz window. The corresponding 2D *D*_iso_-*D*_Δ_^2^ projection comprises a single peak at E[*D*_iso_] = 0.2·10^−9^ m^2^s^−1^ and E[*D*_Δ_^2^] moving slightly from 0.89 at 35 Hz to 0.85 at 320 Hz. The presence of just one peak shows that exchange averaging over the ~50 ms duration of the gradient waveform have rendered the detectable water populations too similar to resolve within the variability of the 2D *D*_iso_-*D*_Δ_^2^ projections of the 100 replicate solutions obtained by bootstrapping and Monte Carlo data inversion. According to Eq. (15), the value of *D*_Δ_^2^ reaches unity in the extreme case of *D*_A_ >> *D*_R_ and *D*_R_ = 0, corresponding to one-dimensional diffusion in an infinitesimally thin cylinder. The observed values of E[*D*_Δ_^2^] are close to one-dimensional diffusion, but still accommodate sufficient displacements in the radial directions to mix water populations in the intraaxonal and adjacent extracellular spaces via the gaps between the myelin patches or directly across the sheaths (Le Bihan et al., 1993). Alternatively, the observations are consistent also with a scenario where the populations remain separate but coincidentally have too similar values of both *D*_iso_ and *D*_Δ_^2^ to resolve without postulating their existence as in the popular model-based approaches (Assaf and Basser, 2005; Zhang et al., 2012).

GM comprises mainly neuronal cell bodies, glial cells, and nonmyelinated axons with low orientational order. The cortex and cerebellum GM data in Figure 2e and f resemble the tumor data in Figure 2c, with negligible influence of Θ and Φ at constant *b, τ*_R_, and *τ*_E_, indicating isotropic diffusion. The deviations between the residuals from the *ω*-dependent and *ω*-independent analyses are clearly visible for both the cortex and cerebellum, but the magnitude is larger for the latter case indicating more pronounced *ω*-dependence in the ~30-300 Hz range. Both examples feature single peaks in the 2D *D*_iso_-*D*_Δ_^2^ projections, with a shift of the peak maximum with frequency being readily apparent for the latter. The explicit shifts are from E[*D*_iso_] = 0.33·10^−9^ m^2^s^−1^ and E[*D*_Δ_^2^] = 0.07 at 35 Hz to E[*D*_iso_] = 0.37·10^−9^ m^2^s^−1^ and E[*D*_Δ_^2^] = 0.05 at 320 Hz for the cortex and from E[*D*_iso_] = 0.28·10^−9^ m^2^s^−1^ and E[*D*_Δ_^2^] = 0.16 at 35 Hz to E[*D*_iso_] = 0.58·10^−9^ m^2^s^−1^ and E[*D*_Δ_^2^] = 0.03 at 320 Hz for the cerebellum. The *ω*-dependence of E[*D*_iso_] can be reproduced with the closed spherical compartment model (Stepišnik, 1993) using 0.4·10^−9^ m^2^s^−1^ local diffusivity and 4 μm radius for the cortex and 0.7·10^−9^ m^2^s^−1^ local diffusivity and 5 μm radius for the cerebellum. Among many other possible mechanisms, these lower and higher values of the local diffusivity could result from the biologically plausible macromolecular contents of, respectively, 30 and 10 vol% (Topgaard, 2020) in solutions with salt and metabolites reducing the diffusivity to 50% of the pure water reference state. As stated above, this model-based interpretation is certainly oversimplified but here serves the purpose to illustrate that rather subtle differences in local chemical composition or biomembrane geometry may have a large impact on data acquired under conditions that are determined more by hardware constraints than by the wishes of the experimentalist.

If **D**(*ω*) depends on *ω* within the investigated window, the *ω*-dependent and *ω*-independent analyses give different residuals as well as 2D *D*_iso_-*D*_Δ_^2^ projections. The difference for the latter is minimized if the *ω*-dependent projection is evaluated at the center of the investigated frequency range as quantified by the *ω*_50%_ metric shown in Figure 1f. Visual inspection of the residuals and 2D *D*_iso_-*D*_Δ_^2^ projections in Figure 2 reveals a correlation between misfit and bias towards higher values of *D*_Δ_^2^ with only minor influence on *D*_iso_. The magnitude of the bias is investigated further in Figure 3 showing parameter maps extracted from the distributions obtained by the *ω*-dependent (top row) and *ω*-independent (middle row) analyses, the former being evaluated at *ω*_50%_ = 190 Hz. The *ω*-dependence as such is reported in terms of the model-independent dispersion metrics Δ_*ω*/2π_E[*D*_iso_] and Δ_*ω*/2π_E[*D*_Δ_^2^] defined in Eq. (20). Superficially, the two analysis approaches appear to give similar parameter maps, except for E[*D*_Δ_^2^] with noticeably lower values in GM for the *ω*-dependent analysis. For the cerebellum, this effect leads to sharper differentiation between GM and WM. The bias in E[*D*_Δ_^2^] is mirrored in the bin-resolved signal fractions map, showing lower values of the anisotropic fraction (bin1) in the *ω*-dependent results. The minor differences between the maps are amplified in the normalized difference maps (bottom row) at the expense of exaggerating the deviations when the metrics are near zero. In general, the difference maps are positive (+10%) for E[*D*_iso_], negative (–100%) for E[*D*_Δ_^2^], and close to zero for *S*_0_, E[*R*_1_], and E[*R*_2_]. An exception to this general observation is the right part of the cerebellum which seems to be contaminated by an imaging artifact affecting primarily *S*_0_ and E[*R*_1_]. While *S*_0_, *R*_1_, and *R*_2_ do not have any explicit *ω*-dependence, a poor fit in the diffusion dimensions could introduce a bias also in the other metrics. The areas highlighted in the Δ_*ω*/2π_E[*D*_iso_] and Δ_*ω*/2π_E[*D*_Δ_^2^] maps, such as the cerebellar GM and the tip of the lateral ventricles (Aggarwal et al., 2012), coincide with the bias in E[*D*_iso_] and E[*D*_Δ_^2^].

**Figure 3:**
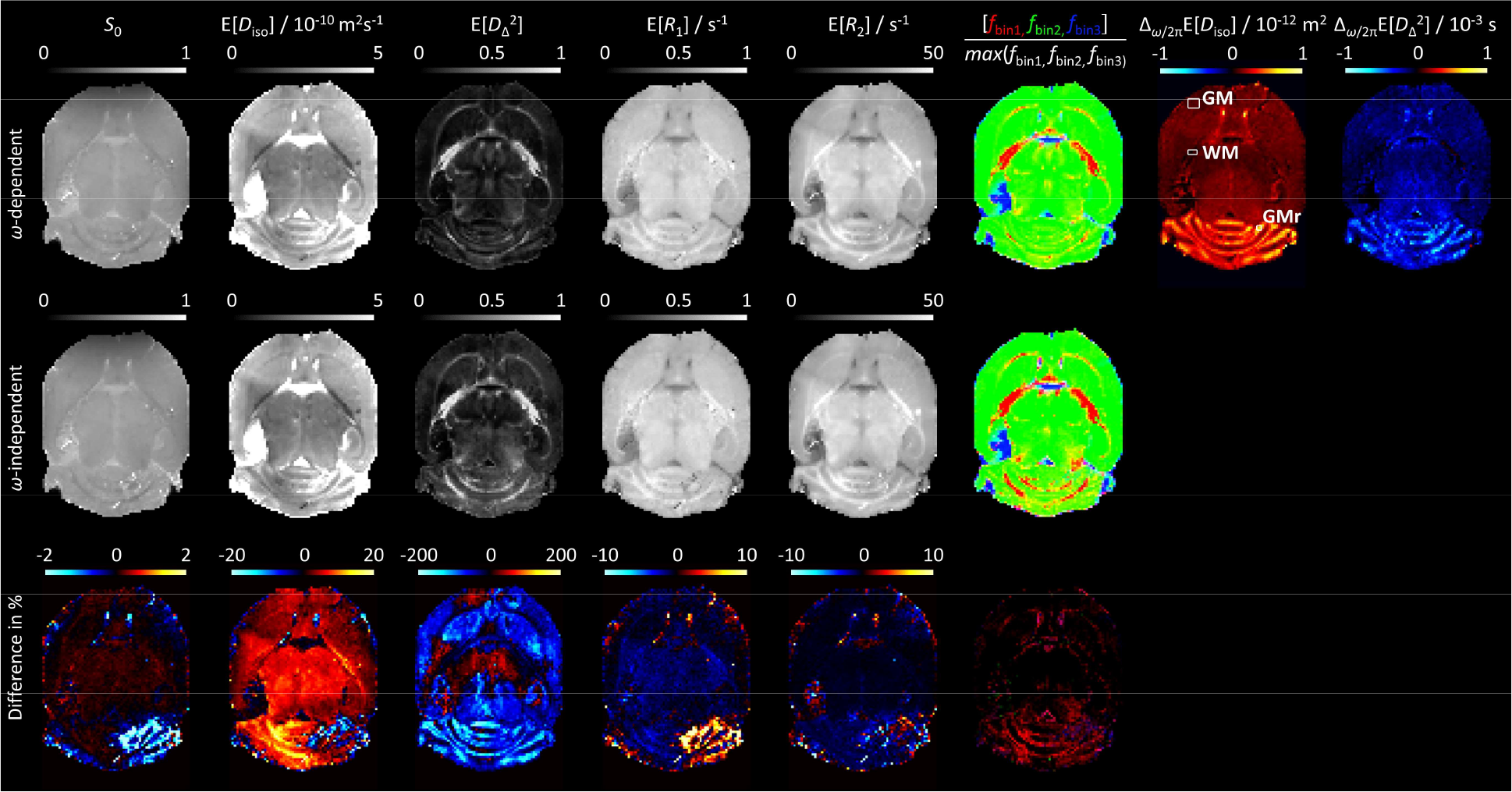
Ex vivo mouse brain parameter maps obtained from *ω*-dependent and *ω*-independent inversions of data acquired with the preclinical (wide *ω*_cent_-range) protocol in Figure 1c. The primary distributions were converted to extrapolated signals *S*_0_, means E[*X*], and bin-resolved signal fractions *f*_bin*n*_ via Eqs. (17), (16), and (19), respectively, using the frequency *ω*_50%_ = 190 Hz labeled in Figure 1f for the *ω*-dependent case. The color-coding of *f*_bin*n*_ is given in Eq. (18). The *ω*-dependence metrics Δ_*ω*/2π_E[*X*], defined in Eq. (20), were evaluated using the frequencies *ω*_10%_ = 35 Hz and *ω*_90%_ = 320 Hz shown in Figure 1f. The normalized differences were calculated according to Eq. (21).

Figure 4 and Figure 5 show results for in vivo human brain obtained with the narrow *ω*_cent_-range protocol in Figure 1e. In addition to the numerous exchange and *ω*-dependence mechanisms described above for water in the salt solution, liquid crystal, tumor tissue, and fixated mouse brain, the living brain features processes originating from the beating heart and the varying pressure in the blood vessels. These processes include not only blood flow in the arteries, veins, and capillary network, as well as pulsatile motion of the entire brain (Wagshul et al., 2011), but also flow of cerebrospinal fluid (CSF) in the ventricles, interstitial fluid (ISF) in the extracellular spaces, and mixed CSF and ISF in the perivascular spaces (Jessen et al., 2015), giving rise to dispersion in **D**(*ω*) at frequencies determined by the interplay between the fluid flow rates and vessel curvatures (Callaghan and Stepišnik, 1995). The gradient waveforms in Figure 1d were designed within hardware constraints with the aim of minimizing *τ*_E_ for given values of *b* and *b*_Δ_, unintentionally giving rise to an anisotropic spread of spectral power in **b**(*ω*) for each waveform (Lundell and Lasič, 2020) as well as a variation of *ω*_cent_ from *ω*_10%_ = 5 Hz to *ω*_90%_ = 11 Hz. Although the relative variation of *ω*_cent_ is sufficient to detect *ω*-dependence as previously demonstrate for ex vivo rat brain (Narvaez et al., 2022), the data for WM, GM, and CSF ROIs in Figure 4 show no clear differences between the residuals or 2D *D*_iso_-*D*_Δ_^2^ projections from the *ω*-dependent and *ω*-independent analyses. This absence of *ω*-dependence in the 5-11 Hz range is far from obvious considering the continuous range of structural length scales and dynamical time scales known to exist in the living brain, but is consistent with literature results of identical diffusion tensor distributions at the diffusion times 19 and 49 ms (Song et al., 2022) and the numerous oscillating gradient spin-echo studies finding dispersion predominantly at higher frequencies (Baron and Beaulieu, 2014; Van et al., 2014; Baron et al., 2015; Arbabi et al., 2020; Tan et al., 2020; Tetreault et al., 2020; Hennel et al., 2021; Michael et al., 2022; Dai et al., 2023).

**Figure 4:**
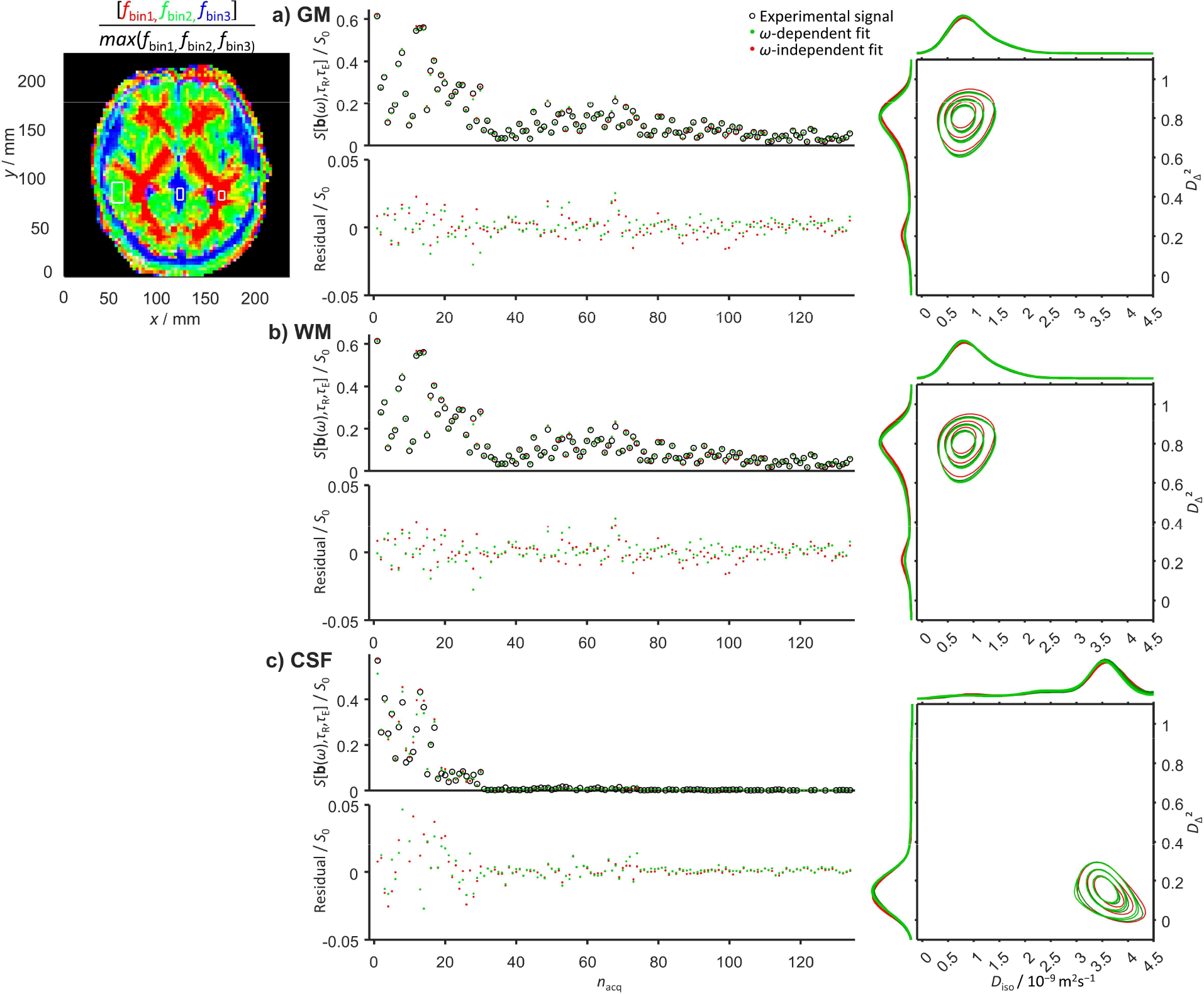
Comparison between *ω*-dependent and *ω*-independent data inversion results for in vivo human brain ROIs in gray matter (GM), white matter (WM), and cerebrospinal fluid (CSF). The data was acquired with the clinical (narrow *ω*_cent_-range) protocol in Figure 1e and the *ω*-dependent 2D *D*_iso_-*D*_Δ_^2^ projections are plotted for three values of *ω* between *ω*_10%_ = 5 Hz (dark green) and *ω*_90%_ = 11 Hz (pale green). For additional explanations of symbols and labels, see Figure 2 caption.

**Figure 5:**
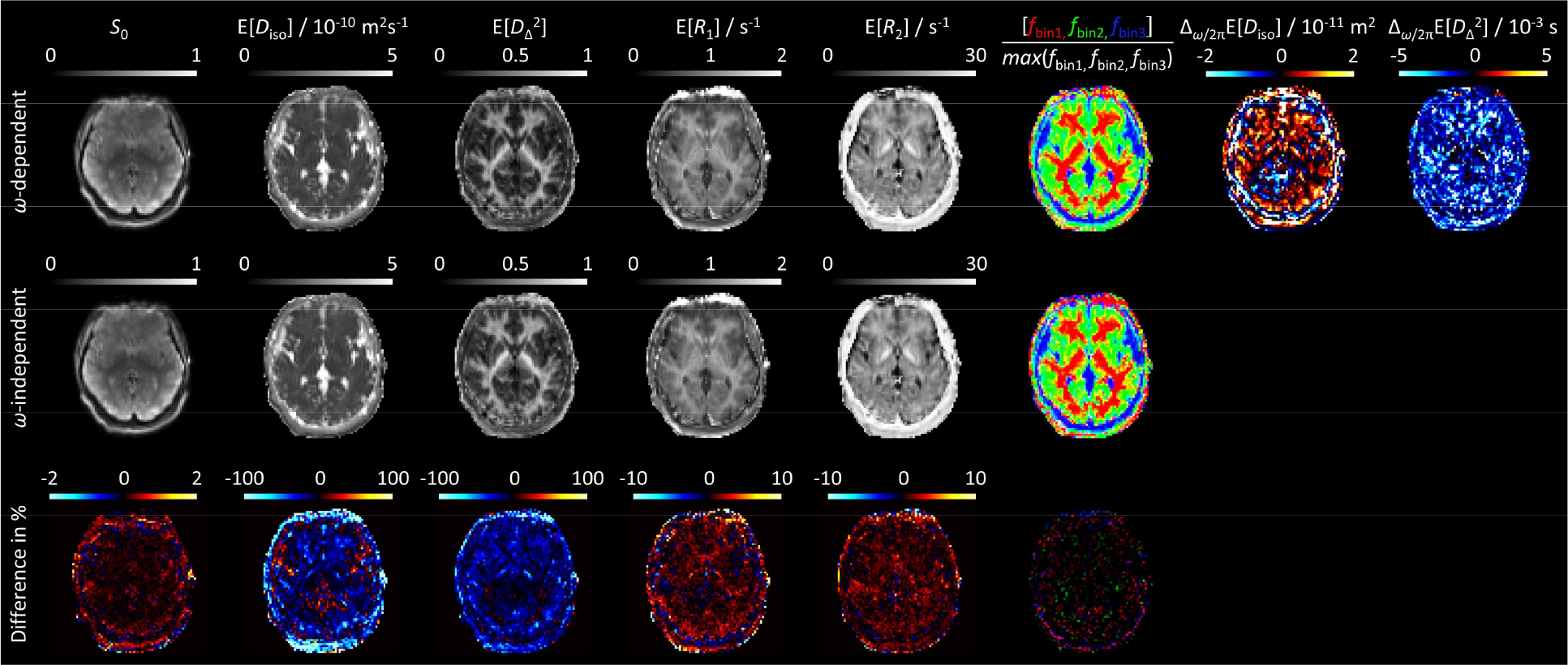
In vivo human brain parameters maps obtained from *ω*-dependent and *ω*-independent inversions of data acquired with the clinical (narrow *ω*-range) protocol in Figure 1e. The *ω*-dependent E[*X*] maps were evaluated at *ω*_50%_ = 9 Hz while the *ω*-dependence metrics Δ_*ω*/2π_E[*X*] employed *ω*_10%_ = 5 Hz and *ω*_90%_ = 11 Hz labeled in Figure 1g. For additional explanations, see Figure 3 caption.

The overall description and interpretation of the in vivo WM and GM data in Figure 4a and b are similar to the ex vivo results above, one minor difference being an oblate component visible in the 1D *D*_Δ_^2^ projection for WM at *D*_Δ_^2^ slightly below 0.25. Spurious oblate components in general appear as inversion artifacts at low signal-to-noise ratio and insufficient exploration of the *b*_Δ_ acquisition dimension (de Almeida Martins and Topgaard, 2018). For acquisition protocols limited to *b*_Δ_ = 1, such oblate components may even dominate the distributions for most voxels except CSF and coherently aligned WM (Song et al., 2022).

The *ω*-dependent and *ω*-independent 1D *D*_Δ_^2^ projections for GM are not completely overlapping despite the absence of discernible differences in the corresponding residuals. This effect may originate from the correlation between *ω*_cent_ and *b*_Δ_ at the highest values of *b* in the acquisition protocol in Figure 1e and the slightly lower signal intensities observed for *ω*_cent_/2π = 10 Hz and *b*_Δ_ = –0.5 than for *ω*_cent_/2π = 6 Hz and *b*_Δ_ = 1. At constant *b*, the powder-averaged signal would increase with *b*_Δ_^2^ and decrease with *ω*_cent_ for the cases of anisotropy and restriction, respectively (Jiang et al., 2023). In modified versions of the acquisition protocol, disambiguation between the two cases could be achieved by extending the *ω*_cent_ range for the two values of *b*_Δ_ at the expense of increasing *τ*_E_. The CSF data in Figure 4c show elevated residuals at low *b* which by visual inspection of the raw images may be attributed to signal dropouts from pulsation artifacts (Chen et al., 2015). Because of the poor fit, it is unclear if the obtained distributions with E[*D*_iso_] = 3.5·10^−9^ m^2^s^−1^, approx. 20% above the literature value 3.0·10^−9^ m^2^s^−1^ for pure water at 37 ºC (Holz et al., 2000), is influenced by intravoxel CSF flow. The absence of *ω*-dependence within the investigated 5-11 Hz range is further accentuated by the similarities of the *ω*-dependent and *ω*-independent parameter maps and the lack of brain-specific structures in the difference and dispersion metrics maps in Figure 5.

## Conclusion

Monte Carlo inversion of multidimensional diffusion-relaxation correlation MRI data can be augmented with explicit inclusion of frequency-dependence of the tensor-valued diffusion spectrum for acquisition protocols exploring both wide and narrow frequency ranges and samples with and without observable effects of restricted diffusion within the investigated frequency window. In the former case, inversion including frequency-dependence gives smaller fit residuals and mitigates a positive bias in anisotropy metrics. In the latter case, inversions with and without consideration of frequency-dependence give similar fit residuals and parameter maps. Consequently, we recommend that frequency-dependence is included by default to avoid bias in the diffusion metrics and allow quantification of the effects of restriction when present.

## Ethics

The research was conducted according to the principles expressed in the Declaration of Helsinki. Participants gave voluntary informed consent for this study. The human and animal studies were approved by the local institutional review boards.

## Data and Code Availability

MATLAB source code for Monte-Carlo data inversion is freely available at https://github.com/daniel-topgaard/md-dmri/.

## Author Contributions

MY: pulse sequence development, data acquisition and processing, manuscript drafting. ON: ex vivo mouse preparation, data acquisition. JM: pulse sequence development, heathy volunteer recruitment, data acquisition and pre-processing. HJ: phantom development and manufacturing, data acquisition. DB: tumor preparation. EFA: tumor model development. FL: pulse sequence development. AS: mouse model development. DT: conceptualization, data processing, manuscript revision. All authors contributed to the final version of the manuscript.

## References

Aggarwal, M., Burnsed, J., Martin, L.J., Northington, F.J., Zhang, J., 2014. Imaging neurodegeneration in the mouse hippocampus after neonatal hypoxia-ischemia using oscillating gradient diffusion MRI. Magn. Reson. Med. 72, 829–840. doi: 10.1002/mrm.24956

Aggarwal, M., Jones, M.V., Calabresi, P.A., Mori, S., Zhang, J., 2012. Probing mouse brain microstructure using oscillating gradient diffusion MRI. Magn. Reson. Med. 67, 98–109. doi: 10.1002/mrm.22981

Aranda, R., Ramirez-Manzanares, A., Rivera, M., 2015. Sparse and adaptive diffusion dictionary (SADD) for recovering intra-voxel white matter structure. Med. Image Anal. 26, 243–255. doi: 10.1016/j.media.2015.10.002

Arbabi, A., Kai, J., Khan, A.R., Baron, C.A., 2020. Diffusion dispersion imaging: Mapping oscillating gradient spin-echo frequency dependence in the human brain. Magn Reson Med 83, 2197–2208. doi: 10.1002/mrm.28083

Assaf, Y., Basser, P.J., 2005. Composite hindered and restricted model of diffusion (CHARMED) MR imaging of the human brain. Neuroimage 27, 48–58. doi: 10.1016/j.neuroimage.2005.03.042

Balucani, U., Brodholt, J.P., Vallauri, R., 1996. Analysis of the velocity autocorrelation function of water. J. Phys.: Condens. Matter 8, 6139–6144. doi: 10.1088/0953-8984/8/34/004

Banani, S.F., Lee, H.O., Hyman, A.A., Rosen, M.K., 2017. Biomolecular condensates: organizers of cellular biochemistry. Nat. Rev. Mol. Cell Biol. 18, 285–298. doi: 10.1038/nrm.2017.7

Baron, C.A., Beaulieu, C., 2014. Oscillating gradient spin-echo (OGSE) diffusion tensor imaging of the human brain. Magn. Reson. Med. 72, 726–736. doi: 10.1002/mrm.24987

Baron, C.A., Kate, M., Gioia, L., Butcher, K., Emery, D., Budde, M., Beaulieu, C., 2015. Reduction of diffusion-weighted imaging contrast of acute ischemic stroke at short diffusion times. Stroke 46, 2136–2141. doi: 10.1161/STROKEAHA.115.008815

Basser, P.J., Mattiello, J., Le Bihan, D., 1994. Estimation of the effective self-diffusion tensor from the NMR spin echo. J. Magn. Reson. B 103, 247–254. doi: 10.1006/jmrb.1994.1037

Basser, P.J., Pajevic, S., 2003. A normal distribution for tensor-valued random variables: applications to diffusion tensor MRI. IEEE Trans Med Imaging 22, 785–794. doi: 10.1109/TMI.2003.815059

Beaulieu, C., 2002. The basis of anisotropic water diffusion in the nervous system - a technical review. NMR Biomed. 15, 435–455. doi: 10.1002/nbm.782

Ben-Shaul, A., Gelbart, W.M., 1985. Theory of chain packing in amphiphilic aggregaes. Ann. Rev. Phys. Chem. 36, 179–211. doi:

Benjamini, D., Basser, P.J., 2020. Multidimensional correlation MRI. NMR Biomed., e4226. doi: 10.1002/nbm.4226

Berman, P., Levi, O., Parmet, Y., Saunders, M., Wiesman, Z., 2013. Laplace inversion of low-resolution NMR relaxometry data using sparse representation methods. Conc. Magn. Reson. A 42, 72–88. doi: 10.1002/cmr.a.21263

Bernin, D., Koch, V., Nydén, M., Topgaard, D., 2014. Multi-scale characterization of lyotropic liquid crystals using ^2^H and diffusion MRI with spatial resolution in three dimensions. PLOS ONE 9, e98752. doi: 10.1371/journal.pone.0098752

Bernin, D., Topgaard, D., 2013. NMR diffusion and relaxation correlation methods: New insights in heterogeneous materials. Curr. Opin. Colloid Interface Sci. 18, 166–172. doi: 10.1016/j.cocis.2013.03.007

Bleuzen, A., Pittet, P.-A., Helm, L., Merbach, A.E., 1997. Water exchange on magnesium(II) in aqueous solution: A variable temperature and pressure ^17^O NMR study. Magn. Reson. Chem. 35, 765–773. doi: 10.1002/(sici)1097-458x(199711)35:11<765::Aid-omr169>3.0.Co;2-f

Boss, B.D., Stejskal, E.O., 1965. Anisotropic diffusion in hydrated vermiculite. J. Chem. Phys. 43, 1068–1069. doi: 10.1063/1.1696823

Callaghan, P.T., Stepišnik, J., 1995. Frequency-domain analysis of spin motion using modulated-gradient NMR. J. Magn. Reson. A 117, 118–122. doi: 10.1006/jmra.1995.9959

Chen, L., Beckett, A., Verma, A., Feinberg, D.A., 2015. Dynamics of respiratory and cardiac CSF motion revealed with real-time simultaneous multi-slice EPI velocity phase contrast imaging. Neuroimage 122, 281–287. doi: 10.1016/j.neuroimage.2015.07.073

Clark, C.A., Hedehus, M., Moseley, M.E., 2001. Diffusion time dependence of the apparent diffusion tensor in healthy human brain and white matter disease. Magn. Reson. Med. 45, 1126–1129. doi: 10.1002/mrm.1149

Colvin, D.C., Loveless, M.E., Does, M.D., Yue, Z., Yankeelov, T.E., Gore, J.C., 2011. Earlier detection of tumor treatment response using magnetic resonance diffusion imaging with oscillating gradients. Magn. Reson. Imaging 29, 315–323. doi: 10.1016/j.mri.2010.10.003

Colvin, D.C., Yankeelov, T.E., Does, M.D., Yue, Z., Quarles, C., Gore, J.C., 2008. New insights into tumor microstructure using temporal diffusion spectroscopy. Cancer Res. 68, 5941–5947. doi: 10.1158/0008-5472.CAN-08-0832

Conturo, T.E., McKinstry, R.C., Akbudak, E., Robinson, B.H., 1996. Encoding of anisotropic diffusion with tetrahedral gradients: A general mathematical diffusion formalism and experimental results. Magn. Reson. Med. 35, 399–412. doi: 10.1002/mrm.1910350319

Cooper, R.L., Chang, D.B., Young, A.C., Martin, C.J., Ancker-Johnson, B., 1974. Restricted diffusion in biophysical systems: Experiment. Biophys J. 14, 161–177. doi: 10.1016/S0006-3495(74)85904-7

Cordero-Grande, L., Christiaens, D., Hutter, J., Price, A.N., Hajnal, J.V., 2019. Complex diffusion-weighted image estimation via matrix recovery under general noise models. Neuroimage 200, 391–404. doi: 10.1016/j.neuroimage.2019.06.039

Dai, E., Zhu, A., Yang, G.K., Quah, K., Tan, E.T., Fiveland, E., Foo, T.K.F., McNab, J.A., 2023. Frequency-dependent diffusion kurtosis imaging in the human brain using an oscillating gradient spin echo sequence and a high-performance head-only gradient. Neuroimage 279, 120328. doi: 10.1016/j.neuroimage.2023.120328

Daimiel Naranjo, I., Reymbaut, A., Brynolfsson, P., Lo Gullo, R., Bryskhe, K., Topgaard, D., Giri, D.D., Reiner, J.S., Thakur, S., Pinker-Domenig, K., 2021. Multidimensional diffusion magnetic resonance imaging for characterization of tissue microstructure in breast cancer patients: A prospective pilot study. Cancers 13, 1606. doi: 10.3390/cancers13071606

de Almeida Martins, J.P., Tax, C.M.W., Reymbaut, A., Szczepankiewicz, F., Chamberland, M., Jones, D.K., Topgaard, D., 2021. Computing and visualising intra-voxel orientation-specific relaxation-diffusion features in the human brain. Hum. Brain Mapp. 42, 310–328. doi: 10.1002/hbm.25224

de Almeida Martins, J.P., Tax, C.M.W., Szczepankiewicz, F., Jones, D.K., Westin, C.-F., Topgaard, D., 2020. Transferring principles of solid-state and Laplace NMR to the field of in vivo brain MRI. Magn. Reson. 1, 27–43. doi: 10.5194/mr-1-27-2020

de Almeida Martins, J.P., Topgaard, D., 2016. Two-dimensional correlation of isotropic and directional diffusion using NMR. Phys. Rev. Lett. 116, 087601. doi: 10.1103/PhysRevLett.116.087601

de Almeida Martins, J.P., Topgaard, D., 2018. Multidimensional correlation of nuclear relaxation rates and diffusion tensors for model-free investigations of heterogeneous anisotropic porous materials. Sci. Rep. 8, 2488. doi: 10.1038/s41598-018-19826-9

de Swiet, T.M., Mitra, P.P., 1996. Possible systematic errors in single-shot measurements of the trace of the diffusion tensor. J. Magn. Reson. B 111, 15–22. doi: 10.1006/jmrb.1996.0055

Does, M.D., Parsons, E.C., Gore, J.C., 2003. Oscillating gradient measurements of water diffusion in normal and globally ischemic rat brain. Magn. Reson. Med. 49, 206–215. doi: 10.1002/mrm.10385

Edelstein, W.A., Hutchison, J.M.S., Johnson, G., Redpath, T., 1980. Spin warp NMR imaging and applications to human whole-body imaging. Phys. Med. Biol. 25, 751–756. doi: 10.1088/0031-9155/25/4/017

Edzes, H.T., Samulski, E.T., 1975. Cross relaxation and spin diffusion in the proton NMR of hydrated collagen. Nature 265, 521–523. doi:

Ekwall, P., Mandell, L., Fontell, K., 1969. Ternary systems of potassium soap, alcohol, and water. J. Colloid Interface Sci. 31, 508–529. doi: 10.1016/0021-9797(69)90052-6

Eriksson, S., Lasič, S., Nilsson, M., Westin, C.-F., Topgaard, D., 2015. NMR diffusion encoding with axial symmetry and variable anisotropy: Distinguishing between prolate and oblate microscopic diffusion tensors with unknown orientation distribution. J. Chem. Phys. 142, 104201. doi: 10.1063/1.4913502

Eriksson, S., Lasič, S., Topgaard, D., 2013. Isotropic diffusion weighting by magic-angle spinning of the q-vector in PGSE NMR. J. Magn. Reson. 226, 13–18. doi: 10.1016/j.jmr.2012.10.015

Evans, D.F., Wennerström, H., 1999. The colloidal domain: Where physics, chemistry, biology, and technology meet, 2nd ed. Wiley-VCH, New York.

Galvosas, P., Callaghan, P.T., 2010. Multi-dimensional inverse Laplace spectroscopy in the NMR of porous media. C. R. Phys. 11, 172–180. doi: 10.1016/j.crhy.2010.06.014

Helm, L., Merbach, A.E., 1999. Water exchange on metal ions: experiments and simulations. Coord. Chem. Rev. 187, 151–181. doi: 10.1016/S0010-8545(99)90232-1

Hennel, F., Michael, E.S., Pruessmann, K.P., 2021. Improved gradient waveforms for oscillating gradient spin-echo (OGSE) diffusion tensor imaging. NMR Biomed 34, e4434. doi: 10.1002/nbm.4434

Hennig, J., Nauerth, A., Friedurg, H., 1986. RARE imaging: a fast imaging method for clinical MR. Magn. Reson. Med. 3, 823–833. doi: 10.1002/mrm.1910030602

Hills, B.P., 1990. Nuclear magnetic resonance relaxation studies of proton exchange in methanol-water mixtures. J. Chem. Soc. Faraday Trans. 86, 481–487. doi: 10.1039/FT9908600481

Hills, B.P., Wright, K.M., Belton, P.S., 1989. Proton N.M.R. studies of chemical and diffusive exchange in carbohydrate systems. Mol. Phys. 67, 1309–1326. doi: 10.1080/00268978900101831

Holz, M., Heil, S.R., Sacco, A., 2000. Temperature-dependent self-diffusion coefficients of water and six selected molecular liquids for calibration in accurate ^1^H NMR PFG measurements. Phys. Chem. Chem. Phys. 2, 4740–4742. doi: 10.1039/b005319h

Hyman, A.A., Weber, C.A., Jülicher, F., 2014. Liquid-liquid phase separation in biology. Annu. Rev. Cell. Dev. Biol. 30, 39–58. doi: 10.1146/annurev-cellbio-100913-013325

Jespersen, S.N., Olesen, J.L., Ianus, A., Shemesh, N., 2019. Effects of nongaussian diffusion on “isotropic diffusion” measurements: An ex-vivo microimaging and simulation study. J. Magn. Reson. 300, 84–94. doi: 10.1016/j.jmr.2019.01.007

Jessen, N.A., Munk, A.S., Lundgaard, I., Nedergaard, M., 2015. The glymphatic system: A beginner’s guide. Neurochem. Res. 40, 2583–2599. doi: 10.1007/s11064-015-1581-6

Jian, B., Vemuri, B.C., Özarslan, E., Carney, P.R., Mareci, T.H., 2007. A novel tensor distribution model for the diffusion-weighted MR signal. Neuroimage 37, 164–176. doi: 10.1016/j.neuroimage.2007.03.074

Jiang, H., de Almeida Martins, J.P., Lundberg, D., Tax, C.M.W., Topgaard, D., 2021. Lamellar liquid crystal phantom for validating MRI methods to distinguish oblate and prolate diffusion tensors on whole-body scanners. Proc. Intl. Soc. Mag. Reson. Med. 29, 3417. doi:

Jiang, H., Svenningsson, L., Topgaard, D., 2023. Multidimensional encoding of restricted and anisotropic diffusion by double rotation of the q vector. Magn. Reson. 4, 73–85. doi: 10.5194/mr-4-73-2023

Johnson Jr, C.S., 1993. Effects of chemical exchange in diffusion-ordered 2D NMR spectra. J. Magn. Reson. A 102, 214–218. doi: 10.1006/jmra.1993.1093

Jones, D.K. (Ed.), 2010. Diffusion MRI: Theory, Methods, and Applications. Oxford University Press.

Jost, W., 1952. Diffusion in Solids, Liquids, and Gases. Academic Press, New York.

Kärger, J., 1969. Zur Bestimmung der Diffusion in einem Zweibereichsystem mit Hilfe von gepulsten Feldgradienten. Ann. Phys. 479, 1–4. doi: 10.1002/andp.19694790102

Kellner, E., Dhital, B., Kiselev, V.G., Reisert, M., 2016. Gibbs-ringing artifact removal based on local subvoxel-shifts. Magn. Reson. Med. 76, 1574–1581. doi: 10.1002/mrm.26054

Kingsley, P.B., 2006. Introduction to diffusion tensor imaging mathematics: Part II. Anisotropy, diffusion-weighting factors, and gradient encoding schemes. Conc. Magn. Reson. A 28A, 123–154. doi: 10.1002/cmr.a.20049

Klein, S., Staring, M., Murphy, K., Viergever, M.A., Pluim, J.P., 2010. elastix: A toolbox for intensity-based medical image registration. IEEE Trans. Med. Imaging 29, 196–205. doi: 10.1109/TMI.2009.2035616

Koay, C.G., Basser, P.J., 2006. Analytically exact correction scheme for signal extraction from noisy magnitude MR signals. J. Magn. Reson. 179, 317–322. doi: 10.1016/j.jmr.2006.01.016

Lasič, S., Szczepankiewicz, F., Eriksson, S., Nilsson, M., Topgaard, D., 2014. Microanisotropy imaging: quantification of microscopic diffusion anisotropy and orientational order parameter by diffusion MRI with magic-angle spinning of the q-vector. Front. Physics 2, 11. doi: 10.3389/fphy.2014.00011

Latour, L.L., Kleinberg, R.L., Mitra, P.P., Sotak, C.H., 1995. Pore-size distributions and tortuosity in heterogeneous porous media. J. Magn. Reson. A 112, 83–91. doi: 10.1006/jmra.1995.1012

Latour, L.L., Mitra, P.P., Kleinberg, R.L., Sotak, C.H., 1993. Time-dependent diffusion coefficient of fluids in porous media as a probe of surface-to-volume ratio. J. Magn. Reson. A 101, 342–346. doi: 10.1006/jmra.1993.1056

Latour, L.L., Svoboda, K., Mitra, P.P., Sotak, C.H., 1994. Time-dependent diffusion of water in a biological model system. Proc. Natl. Acad. Sci. USA 91, 1229–1233. doi: 10.1073/pnas.91.4.1229

Laukien, G., Schlüter, J., 1956. Impulstechnische Messungen der Spin-Gitter und der Spin-Spin-Relaxationszeiten von Protonen in wäßrigen Lösungen paramagnetischer Ionen. Z. Physik 146, 113–126. doi: 10.1007/BF01326008

Le Bihan, D., Breton, E., Lallemand, D., Grenier, P., Cabanis, E., Laval-Jeantet, M., 1986. MR imaging of intravoxel incoherent motions - application to diffusion and perfusion in neurological disorders. Radiology 161, 401–407. doi: 10.1148/radiology.161.2.3763909

Le Bihan, D., Turner, R., Douek, P., 1993. Is water diffusion restricted in human brain white matter? Neuroreport 4, 887–890. doi: 10.1097/00001756-199307000-00012

Leow, A.D., Zhu, S., Zhan, L., McMahon, K., de Zubicaray, G.I., Meredith, M., Wright, M.J., Toga, A.W., Thompson, P.M., 2009. The tensor distribution function. Magn. Reson. Med. 61, 205–214. doi: 10.1002/mrm.21852

Li, H., Gore, J.C., Xu, J., 2014. Fast and robust measurement of microstructural dimensions using temporal diffusion spectroscopy. J. Magn. Reson. 242, 4–9. doi: 10.1016/j.jmr.2014.02.007

Ligneul, C., Valette, J., 2017. Probing metabolite diffusion at ultra-short time scales in the mouse brain using optimized oscillating gradients and “short”-echo-time diffusion-weighted MRS. NMR Biomed. 30, e3671. doi: 10.1002/nbm.3671

Lindahl, E., Edholm, O., 2000. Mesoscopic undulations and thickness fluctuations in lipid bilayers from molecular dynamics simulations. Biophys. J. 79, 426–433. doi: 10.1016/S0006-3495(00)76304-1

Lundell, H., Lasič, S., 2020. Diffusion encoding with general gradient waveforms. In: Topgaard, D. (Ed.), Advanced Diffusion Encoding Methods in MRI. Royal Society of Chemistry, Cambridge, UK, pp. 12–67. doi: 10.1039/9781788019910-00012

Lundell, H., Nilsson, M., Dyrby, T.B., Parker, G.J.M., Cristinacce, P.L.H., Zhou, F.L., Topgaard, D., Lasič, S., 2019. Multidimensional diffusion MRI with spectrally modulated gradients reveals unprecedented microstructural detail. Sci. Rep. 9, 9026. doi: 10.1038/s41598-019-45235-7

Lundell, H., Sønderby, C.K., Dyrby, T.B., 2015. Diffusion weighted imaging with circularly polarized oscillating gradients. Magn. Reson. Med. 73, 1171–1176. doi: 10.1002/mrm.25211

Lutti, A., Callaghan, P.T., 2007. Measurement of multilamellar onion dimensions under shear using frequency domain pulsed gradient NMR. J. Magn. Reson. 187, 251–257. doi: 10.1016/j.jmr.2007.05.003

MacGregor, R.P., Peemoeller, H., Schneider, M.H., Sharp, A.R., 1983. Anisotropic diffusion of water in wood. J. Appl. Polym. Sci.: Applied Polymer Symposium 37, 901–909. doi:

Magdoom, K.N., Pajevic, S., Dario, G., Basser, P.J., 2021. A new framework for MR diffusion tensor distribution. Sci. Rep. 11, 2766. doi: 10.1038/s41598-021-81264-x

Martin, J., Endt, S., Wetscherek, A., Kuder, T.A., Doerfler, A., Uder, M., Hensel, B., Laun, F.B., 2020. Contrast-to-noise ratio analysis of microscopic diffusion anisotropy indices in q-space trajectory imaging. Z. Med. Phys. 30, 4–16. doi: 10.1016/j.zemedi.2019.01.003

Martin, J., Reymbaut, A., Schmidt, M., Doerfler, A., Uder, M., Laun, F.B., Topgaard, D., 2021. Nonparametric D-R1-R2 distribution MRI of the living human brain. Neuroimage 245, 118753. doi: 10.1016/j.neuroimage.2021.118753

Michael, E.S., Hennel, F., Pruessmann, K.P., 2022. Evaluating diffusion dispersion across an extended range of b-values and frequencies: Exploiting gap-filled OGSE shapes, strong gradients, and spiral readouts. Magn. Reson. Med. 87, 2710–2723. doi: 10.1002/mrm.29161

Mills, R., 1973. Self-diffusion in normal and heavy water in the range 1-45°. J. Phys. Chem. 77, 685–688. doi: 10.1021/j100624a025

Mori, S., van Zijl, P.C.M., 1995. Diffusion weighting by the trace of the diffusion tensor within a single scan. Magn. Reson. Med. 33, 41–52. doi: 10.1002/mrm.1910330107

Morris, K.F., Johnson Jr, C.S., 1992. Diffusion-ordered two-dimensional nuclear magnetic resonance spectroscopy. J. Am. Chem. Soc. 114, 3139–3141. doi: 10.1021/ja00034a071

Moseley, M.E., Kucharczyk, J., Asgari, H.S., Norman, D., 1991. Anisotropy in diffusion-weighted MRI. Magn. Reson. Med. 19, 321–326. doi: 10.1002/mrm.1910190222

Narvaez, O., Svenningsson, L., Yon, M., Sierra, A., Topgaard, D., 2022. Massively multidimensional diffusion-relaxation correlation MRI. Front. Phys. 9, 793966. doi: 10.3389/fphy.2021.793966

Narvaez, O., Yon, M., Jiang, H., Bernin, D., Forssell-Aronsson, E., Sierra, A., Topgaard, D., 2021. Model-free approach to the interpretation of restricted and anisotropic self-diffusion in magnetic resonance of biological tissues. arXiv:2111.07827. doi: 10.48550/arXiv.2111.07827

Neely, J., Connick, R., 1970. Rate of water exchange from hydrated magnesium ion. J. Am. Chem. Soc. 92, 3476–3478. doi: 10.1021/ja00714a048

Nielsen, J.S., Dyrby, T.B., Lundell, H., 2018. Magnetic resonance temporal diffusion tensor spectroscopy of disordered anisotropic tissue. Sci. Rep. 8, 2930. doi: 10.1038/s41598-018-19475-y

Nilsson, M., Lätt, J., Nordh, E., Wirestam, R., Ståhlberg, F., 2009. On the effects of varied diffusion time in vivo: is the diffusion in white matter restricted? Magn. Reson. Imaging 27, 176–187. doi: 10.1016/j.mri.2008.06.003

Nilsson, M., Szczepankiewicz, F., Lampinen, B., Ahlgren, A., de Almeida Martins, J.P., Lasič, S., Westin, C.-F., Topgaard, D., 2018. An open-source framework for analysis of multidimensional diffusion MRI data implemented in MATLAB. Proc. Intl. Soc. Mag. Reson. Med. 26, 5355. doi:

Nilsson, M., Szczepankiewicz, F., van Westen, D., Hansson, O., 2015. Extrapolation-based references improve motion and eddy-current correction of high b-value DWI data: Application in Parkinson’s disease dementia. PLoS One 10, e0141825. doi: 10.1371/journal.pone.0141825

Packer, K.J., Rees, C., 1972. Pulsed NMR studies of restricted diffusion. I. Droplet size distributions in emulsions. J. Colloid Interface Sci. 40, 206–218. doi: 10.1016/0021-9797(72)90010-0

Parsons, E.C., Does, M.D., Gore, J.C., 2003. Modified oscillating gradient pulses for direct sampling of the diffusion spectrum suitable for imaging sequences. Magn. Reson. Imaging 21, 279–285. doi: 10.1016/s0730-725x(03)00155-3

Parsons Jr, E.C., Does, M.D., Gore, J.C., 2006. Temporal diffusion spectroscopy: Theory and implementation in restricted systems using oscillating gradients. Magn. Reson. Med. 55, 75–84. doi: 10.1002/mrm.20732

Persson, E., Halle, B., 2008. Cell water dynamics on multiple time scales. Proc. Natl. Acad. Sci. USA 105, 6266–6271. doi: 10.1073/pnas.0709585105

Portnoy, S., Flint, J.J., Blackband, S.J., Stanisz, G.J., 2013. Oscillating and pulsed gradient diffusion magnetic resonance microscopy over an extended b-value range: Implications for the characterization of tissue microstructure. Magn. Reson. Med. 69, 1131–1145. doi: 10.1002/mrm.24325

Provencher, S.W., 1982. A constrained regularization method for inverting data represented by linear algebraic or integral equations. Computer Phys. Comm. 27, 213–227. doi: 10.1016/0010-4655(82)90173-4

Reymbaut, A., Critchley, J., Durighel, G., Sprenger, T., Sughrue, M., Bryskhe, K., Topgaard, D., 2021. Towards non-parametric diffusion-T1 characterization of crossing fibers in the human brain. Magn. Reson. Med. 85, 2815–2827. doi: 10.1002/mrm.28604

Reymbaut, A., Mezzani, P., de Almeida Martins, J.P., Topgaard, D., 2020a. Accuracy and precision of statistical descriptors obtained from multidimensional diffusion signal inversion algorithms. NMR Biomed. 33, e4267. doi: 10.1002/nbm.4267

Reymbaut, A., Zheng, Y., Li, S., Sun, W., Xu, H., Daimiel Naranjo, I., Thakur, S., Pinker-Domenig, K., Rajan, S., Vanugopal, V.K., Mahajan, V., Mahajan, H., Critchley, J., Durighel, G., Sughrue, M., Bryskhe, K., Topgaard, D., 2020b. Clinical research with advanced diffusion encoding methods in MRI. In: Topgaard, D. (Ed.), Advanced Diffusion Encoding Methods in MRI. Royal Society of Chemistry, Cambridge, UK, pp. 406–429. doi: 10.1039/9781788019910-00406

Reynaud, O., Winters, K.V., Hoang, D.M., Wadghiri, Y.Z., Novikov, D.S., Kim, S.G., 2015. Surface-to-volume ratio mapping of tumor microstructure using oscillating gradient diffusion weighted imaging. Magn. Reson. Med. doi: 10.1002/mrm.25865

Reynaud, O., Winters, K.V., Hoang, D.M., Wadghiri, Y.Z., Novikov, D.S., Kim, S.G., 2016. Pulsed and oscillating gradient MRI for assessment of cell size and extracellular space (POMACE) in mouse gliomas. NMR Biomed. 29, 1350–1363. doi: 10.1002/nbm.3577

Rosenberg, J.T., Grant, S.C., Topgaard, D., 2022. Nonparametric 5D D-R2 distribution imaging with single-shot EPI at 21.1 T: Initial results for in vivo rat brain. J. Magn. Reson. 341, 107256. doi: 10.1016/j.jmr.2022.107256

Rumble, J. (Ed.), 2021. CRC Handbook of Chemistry and Physics, 102 ed. CRC Press, Boca Raton.

Saliani, A., Perraud, B., Duval, T., Stikov, N., Rossignol, S., Cohen-Adad, J., 2017. Axon and myelin morphology in animal and human spinal cord. Front. Neuroanat. 11, 129. doi: 10.3389/fnana.2017.00129

Schachter, M., Does, M.D., Anderson, A.W., Gore, J.C., 2000. Measurements of restricted diffusion using an oscillating gradient spin-echo sequence. J. Magn. Reson. 147, 232–237. doi: 10.1006/jmre.2000.2203

Simoes, M.C., Hughes, K.J., Ingham, D.B., Ma, L., Pourkashanian, M., 2017. Estimation of the thermochemical radii and ionic volumes of complex ions. Inorg. Chem. 56, 7566–7573. doi: 10.1021/acs.inorgchem.7b01205

Sjölund, J., Szczepankiewicz, F., Nilsson, M., Topgaard, D., Westin, C.-F., Knutsson, H., 2015. Constrained optimization of gradient waveforms for generalized diffusion encoding. J. Magn. Reson. 261, 157–168. doi: 10.1016/j.jmr.2015.10.012

Slator, P.J., Palombo, M., Miller, K.L., Westin, C.F., Laun, F., Kim, D., Haldar, J.P., Benjamini, D., Lemberskiy, G., de Almeida Martins, J.P., Hutter, J., 2021. Combined diffusion-relaxometry microstructure imaging: Current status and future prospects. Magn. Reson. Med. 86, 2987–3011. doi: 10.1002/mrm.28963

Song, Y., Ly, I., Fan, Q., Nummenmaa, A., Martinez-Lage, M., Curry, W.T., Dietrich, J., Forst, D.A., Rosen, B.R., Huang, S.Y., Gerstner, E.R., 2022. Measurement of full diffusion tensor distribution using high-gradient diffusion MRI and applications in diffuse gliomas. Front. Phys. 10, 813475. doi: 10.3389/fphy.2022.813475

Song, Y.-Q., Venkataramanan, L., Kausik, R., Heaton, N., 2016. Two-dimensional NMR of diffusion and relaxation. In: Valiullin, R. (Ed.), Diffusion NMR of Confined Systems: Fluid Transport in Porous Solids and Heterogeneous Materials. Royal Society of Chemistry, Cambridge, UK, pp. 111–155. doi: 10.1039/9781782623779-00111

Stejskal, E.O., 1965. Use of spin echoes in a pulsed magnetic-field gradient to study anisotropic, restricted diffusion and flow. J. Chem. Phys. 43, 3597–3603. doi: 10.1063/1.1696526

Stejskal, E.O., Tanner, J.E., 1965. Spin diffusion measurements: Spin echoes in the presence of a time-dependent field gradient. J. Chem. Phys. 42, 288–292. doi: 10.1063/1.1695690

Stepišnik, J., 1981. Analysis of NMR self-diffusion measurements by a density matrix calculation. Physica B 104, 305–364. doi: 10.1016/0378-4363(81)90182-0

Stepišnik, J., 1993. Time-dependent self-diffusion by NMR spin-echo. Physica B 183, 343–350. doi: 10.1016/0921-4526(93)90124-O

Stepišnik, J., Callaghan, P.T., 2000. The long time tail of molecular velocity correlation in a confined fluid: observation by modulated gradient spin-echo NMR. Physica B 292, 296–301. doi: 10.1016/S0921-4526(00)00469-5

Stilbs, P., 1987. Fourier transform pulsed-gradient spin-echo studies of molecular diffusion. Prog. Nucl. Magn. Reson. Spectrosc. 19, 1–45. doi:

Szczepankiewicz, F., Lasič, S., van Westen, D., Sundgren, P.C., Englund, E., Westin, C.-F., Ståhlberg, F., Lätt, J., Topgaard, D., Nilsson, M., 2015. Quantification of microscopic diffusion anisotropy disentangles effects of orientation dispersion from microstructure: Applications in healthy volunteers and in brain tumors. Neuroimage 104, 241–252. doi: 10.1016/j.neuroimage.2014.09.057

Tan, E.T., Shih, R.Y., Mitra, J., Sprenger, T., Hua, Y., Bhushan, C., Bernstein, M.A., McNab, J.A., DeMarco, J.K., Ho, V.B., Foo, T.K.F., 2020. Oscillating diffusion-encoding with a high gradient-amplitude and high slew-rate head-only gradient for human brain imaging. Magn. Reson. Med. 84, 950–965. doi: 10.1002/mrm.28180

Tanner, J.E., 1979. Self diffusion of water in frog muscle. Biophys. J. 28, 107–116. doi: 10.1016/S0006-3495(79)85162-0

Tax, C.M.W., 2020. Estimating chemical and microstructural heterogeneity by correlating relaxation and diffusion. In: Topgaard, D. (Ed.), Advanced Diffusion Encoding Methods in MRI. Royal Society of Chemistry, Cambridge, UK, pp. 186–227. doi: 10.1039/9781788019910-00186

Tetreault, P., Harkins, K.D., Baron, C.A., Stobbe, R., Does, M.D., Beaulieu, C., 2020. Diffusion time dependency along the human corpus callosum and exploration of age and sex differences as assessed by oscillating gradient spin-echo diffusion tensor imaging. Neuroimage 210, 116533. doi: 10.1016/j.neuroimage.2020.116533

Tofts, P., 2003. Quantitative MRI of the brain: Measuring changes caused by disease. John Wiley.

Topgaard, D., 2016a. Director orientations in lyotropic liquid crystals: Diffusion MRI mapping of the Saupe order tensor. Phys. Chem. Chem. Phys. 18, 8545–8553. doi: 10.1039/c5cp07251d

Topgaard, D., 2016b. NMR methods for studying microscopic diffusion anisotropy. In: Valiullin, R. (Ed.), Diffusion NMR of Confined Systems: Fluid Transport in Porous Solids and Heterogeneous Materials. Royal Society of Chemistry, Cambridge, UK, pp. 226–259. doi: 10.1039/9781782623779-00226

Topgaard, D., 2017. Multidimensional diffusion MRI. J. Magn. Reson. 275, 98–113. doi: 10.1016/j.jmr.2016.12.007

Topgaard, D., 2019a. Diffusion tensor distribution imaging. NMR Biomed. 32, e4066. doi: 10.1002/nbm.4066

Topgaard, D., 2019b. Multiple dimensions for random walks. J. Magn. Reson. 306, 150–154. doi: 10.1016/j.jmr.2019.07.024

Topgaard, D., 2020. Translational motion of water in biological tissues – a brief primer. In: Topgaard, D. (Ed.), Advanced Diffusion Encoding Methods in MRI. Royal Society of Chemistry, Cambridge, UK, pp. 1–11. doi: 10.1039/9781788019910-00001

Topgaard, D., Malmborg, C., Söderman, O., 2002. Restricted self-diffusion of water in a highly concentrated W/O emulsion studied using modulated gradient spin-echo NMR. J. Magn. Reson. 156, 195–201. doi: 10.1006/jmre.2002.2556

Tournier, J.D., Smith, R., Raffelt, D., Tabbara, R., Dhollander, T., Pietsch, M., Christiaens, D., Jeurissen, B., Yeh, C.H., Connelly, A., 2019. MRtrix3: A fast, flexible and open software framework for medical image processing and visualisation. Neuroimage 202, 116137. doi: 10.1016/j.neuroimage.2019.116137

Van, A.T., Holdsworth, S.J., Bammer, R., 2014. In vivo investigation of restricted diffusion in the human brain with optimized oscillating diffusion gradient encoding. Magn. Reson. Med. 71, 83–94. doi: 10.1002/mrm.24632

Wadsö, L., Anderberg, A., Åslund, I., Söderman, O., 2009. An improved method to validate the relative humidity generation in sorption balances. Eur. J. Pharm. Biopharm. 72, 99–104. doi: 10.1016/j.ejpb.2008.10.013

Wagshul, M.E., Eide, P.K., Madsen, J.R., 2011. The pulsating brain: A review of experimental and clinical studies of intracranial pulsatility. Fluids Barriers CNS 8, 5. doi: 10.1186/2045-8118-8-5

Wennerström, H., 1973. Proton nuclear magnetic resonance lineshapes in lamellar liquid crystals. Chem. Phys. Lett. 18, 41–44. doi: 10.1016/0009-2614(73)80333-1

Westin, C.-F., Knutsson, H., Pasternak, O., Szczepankiewicz, F., Özarslan, E., van Westen, D., Mattisson, C., Bogren, M., O’Donnell, L., Kubicki, M., Topgaard, D., Nilsson, M., 2016. Q-space trajectory imaging for multidimensional diffusion MRI of the human brain. Neuroimage 135, 345–362. doi: 10.1016/j.neuroimage.2016.02.039

Westin, C.-F., Szczepankiewicz, F., Pasternak, O., Özarslan, E., Topgaard, D., Knutsson, H., Nilsson, M., 2014. Measurement tensors in diffusion MRI: Generalizing the concept of diffusion encoding. Med. Image Comput. Comput. Assist. Interv. 17, 209–216. doi: 10.1007/978-3-319-10443-0_27

Wetscherek, A., Stieltjes, B., Laun, F.B., 2015. Flow-compensated intravoxel incoherent motion diffusion imaging. Magn. Reson. Med. 74, 410–419. doi: 10.1002/mrm.25410

Whittal, K.P., MacKay, A.L., 1989. Quantitative interpretation of NMR relaxation data. J. Magn. Reson. 84, 134–152. doi: 10.1016/0022-2364(89)90011-5

Woessner, D.E., 1963. N.M.R. spin-echo self-diffusion measurements on fluids undergoing restricted diffusion. J. Phys. Chem. 67, 1365–1367. doi: 10.1021/j100800a509

Wu, D., Martin, L.J., Northington, F.J., Zhang, J., 2018. Oscillating-gradient diffusion magnetic resonance imaging detects acute subcellular structural changes in the mouse forebrain after neonatal hypoxia-ischemia. J. Cereb. Blood Flow. Metab., 271678X18759859. doi: 10.1177/0271678X18759859

Xu, J., Li, K., Smith, R.A., Waterton, J.C., Zhao, P., Chen, H., Does, M.D., Manning, H.C., Gore, J.C., 2012. Characterizing tumor response to chemotherapy at various length scales using temporal diffusion spectroscopy. PLoS One 7, e41714. doi: 10.1371/journal.pone.0041714

Yon, M., de Almeida Martins, J.P., Bao, Q., Budde, M.D., Frydman, L., Topgaard, D., 2020. Diffusion tensor distribution imaging of an in vivo mouse brain at ultra-high magnetic field by spatiotemporal encoding. NMR Biomed. 33, e4355. doi: 10.1002/nbm.4355

Zhang, H., Schneider, T., Wheeler-Kingshott, C.A., Alexander, D.C., 2012. NODDI: Practical in vivo neurite orientation dispersion and density imaging of the human brain. Neuroimage 61, 1000–1016. doi: 10.1016/j.neuroimage.2012.03.072

Zimmerman, J.R., 1954. Proton relaxation in Mn++ aqueous solutions. J. Chem. Phys. 22, 950–950. doi: 10.1063/1.1740232

